# Immune gene expression profiling reveals heterogeneity in luminal breast tumors

**DOI:** 10.1101/515486

**Authors:** Bin Zhu, Shelly Lap Ah Tse, Difei Wang, Hela Koka, Tongwu Zhang, Mustapha Abubakar, Priscilla Lee, Feng Wang, Cherry Wu, Koon Ho Tsang, Wing-cheong Chan, Sze Hong Law, Mengjie Li, Wentao Li, Suyang Wu, Zhiguang Liu, Bixia Huang, Han Zhang, Eric Tang, Zhengyan Kan, Soohyeon Lee, Yeon Hee Park, Seok Jin Nam, Mingyi Wang, Xuezheng Sun, Kristine Jones, Bin Zhu, Amy Hutchinson, Belynda Hicks, Ludmila Prokunina-Olsson, Jianxin Shi, Montserrat Garcia-Closas, Stephen Chanock, Xiaohong R. Yang

## Abstract

Disease heterogeneity of immune gene expression patterns of luminal breast cancer (BC) has not been well studied. We performed immune gene expression profiling of tumor and adjacent normal tissue in 92 Asian luminal BC patients and identified three distinct immune subtypes. Tumors in one subtype exhibited signs of T-cell activation, lower *ESR1/ESR2* expression ratio and higher expression of immune checkpoint genes, nonsynonymous mutation burden, *APOBEC*-signature mutations, and increasing body mass index compared to other luminal tumors. Tumors in a second subtype were characterized by increased expression of interferon-stimulated genes and enrichment for *TP53* somatic mutations. The presence of three immune subtypes within luminal BC was replicated in cases drawn from The Cancer Genome Atlas and a Korean breast cancer study. Our findings suggest that immune gene expression and associated genomic features could be useful to further stratify luminal BC beyond the current luminal A/B classification.

## Introduction

Breast cancer (BC) is a heterogeneous disease comprised of several molecular subtypes (luminal A, luminal B, HER2-enriched, and basal-like) with distinct molecular features and clinical behaviors^1,2^. Within each subtype, substantial heterogeneity still exists in terms of genomic features and clinical outcomes^3–5^. Luminal BC is the predominant subtype; currently we cannot precisely identify patients who do not respond to endocrine therapy and carry a poor prognosis^6^. The commonly used luminal A/B classification based on proliferation does not fully capture heterogeneity in luminal tumors^7,8^. A recent study^9^ partitioned luminal breast tumors of The Cancer Genome Atlas (TCGA) into two distinct prognostic subgroups that exhibited differential expression of immune-related genes. This partition showed better discriminative prognostic value than the luminal A/B classification, suggesting that the immunogenicity of luminal tumors is heterogeneous.

The investigation of tumor-infiltrating lymphocytes (TILs) has greatly improved our knowledge of the nature of tumor-immune interactions. The presence of TILs has been associated with a favorable prognosis across multiple cancer types including BC, although TILs might be associated with treatment responses and survival in a subtype-specific manner^10,11,12^, suggesting a dependence of the immune infiltration on BC subtypes. Recent TCGA Pan-Cancer studies identified substantial heterogeneity in immune profiles across and within cancer types as well as within cancer subtypes^13,14^. For example, Thorsson *et al.*^13^identified perhaps six immune subtypes spanning multiple cancer types and most breast tumors fell into three of these immune subtypes. Among BC molecular subtypes, luminal-A tumors showed the greatest heterogeneity, with a similar number of tumors classified into each of the three immune subtypes. Nevertheless, variation in immune profiles within luminal tumors may not be sufficiently characterized in these Pan-Cancer analyses. In analyses including all BC subtypes, the immune stratification was likely driven by HER2-enriched and basal-like tumors since TILs are more abundant in these subtypes than in luminal BC^15^. A more detailed understanding of the variation in TILs among luminal tumors could provide new insights into luminal BC heterogeneity and identify a subset who might be amenable to immunomodulation and benefit from immunotherapy.

So far, most studies that conduct profiles of immune cells in BC have used data from TCGA, which does not represent the general patient population, particularly for non-European subjects. Previous studies have shown that tumor immunobiology might vary by race/ethnicity^16,17^ but the contribution of biological factors to racial differences is largely unknown. Different germline genetic architecture may play a role but how germline variants contribute to immune phenotype has not been extensively studied. For example, the germline *APOBEC3B* deletion polymorphism, which is more common in East Asians (31.2%) than in Europeans (9.0%) and West Africans (4.2%) based on HapMap, is not well represented in TCGA. This deletion has been associated with increased BC risk^18^ and immune gene expression^19,20^, suggesting that East Asian BCs may exhibit a distinct immune profile compared to other BC populations. In this study, we profiled immune gene expression in paired tumor/normal luminal breast tissue collected from a hospital-based case-control study of Asian BC patients in Hong Kong (HKBC), for whom extensive clinical and epidemiologic data were collected.

## Results

The analysis included 92 luminal tumors and 56 normal samples (including 56 tumor/normal tissue pairs) with good quality of RNA-Sequencing data (HKBC). The mean age at diagnosis was 58.7 years; the majority of these patients were postmenopausal (76.1%). 49 (53.3%) and 43 (46.7%) patients were classified as luminal-A and luminal-B, respectively, according to PAM50. Although our analyses were focused on luminal patients, we also present data for HER2-enriched and basal-like patients as a comparison group (n=40). The distribution of clinical characteristics and key BC risk factors among these patients is shown in Supplementary Table 1.

### Immune gene expression stratified luminal tumors into three subtypes

We conducted unsupervised consensus clustering of 92 luminal tumors using expression of 130 immune-related genes (within 13 previously reported metagenes)^21^(Supplementary Table 2). The best separation was achieved by dividing the luminal patients into three subtypes (lum1: n=40; lum2: n=36; lum3; n=16; Figure 1a); lum1 and lum3 were enriched with luminal-A tumors and lum2 enriched with luminal-B tumors (Supplementary Table 3). Lum1 expressed low levels of most immune genes (Figure 1b) and therefore was designated as low-TIL. Lum2 had high expression of *STAT1* and other interferon-stimulated genes (ISGs), but low expression of other immune genes (Figure 1b), designated as high-ISG. Lum3 (defined as high-TIL) showed the highest expression level of most immune genes (Figure 1b) such as immune checkpoint genes (e.g. *PD-L1* and *CTLA-4*), chemokine genes and their receptors (e.g. *CXCL9* and *CXCL10*) and effectors (e.g. *GZMK* and *PRF1*) (Supplementary Figure 1), reflecting a T cell-inflamed phenotype. Compared to low-TIL and high-ISG tumors, high-TIL tumors had significantly higher abundance of most immune subpopulations (estimated by MCP-counter, Figure 2a), except for neutrophils and cells of monocytic lineage. The abundance score for each immune subpopulation in high-TIL luminal tumors was comparable to that of HER2-enriched and basal-like tumors (Figure 2a; P values see Supplementary Table 4). Adjusting for tumor purity, which was inferred using ESTIMATE purity score, did not significantly change the results (Supplementary Figure 2). Representative images of immune infiltration in high-TIL tumors are shown in Supplementary Figure 3.

**Figure 1:**
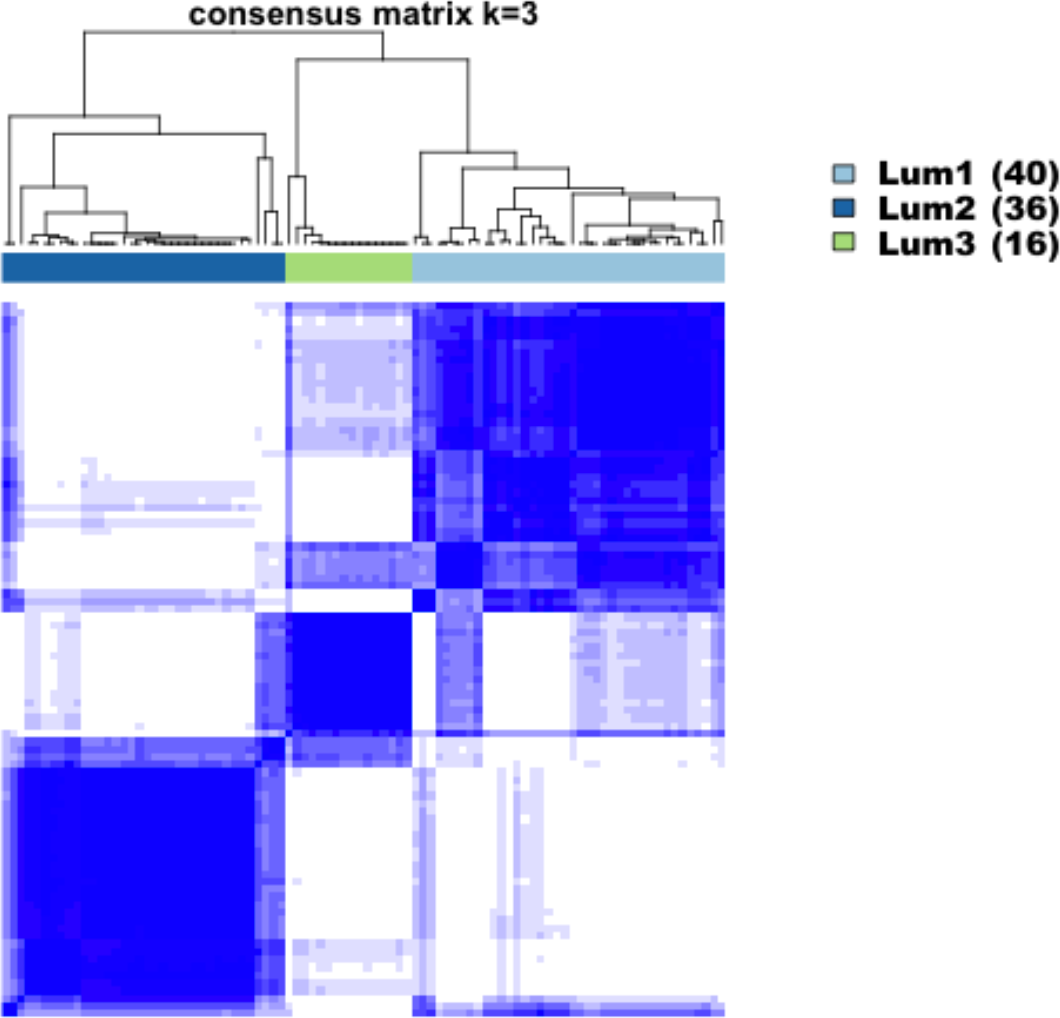
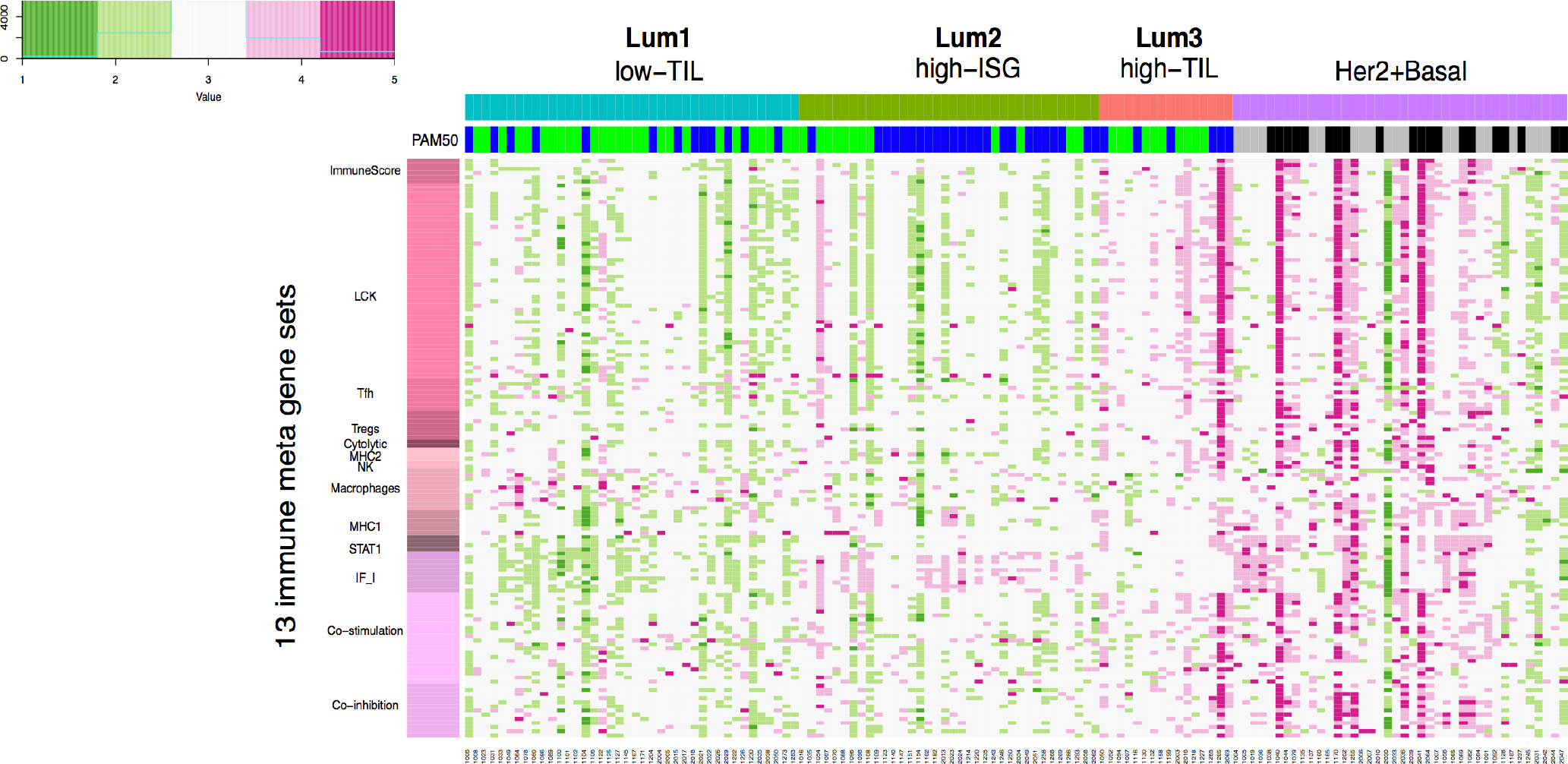
Consensus clustering of 92 luminal breast tumors from Hong Kong patients based on 130 immune-related genes. a) Consensus cluster matrix showing three major clusters; b) Gene expression heatmap showing gene expression levels of 13 immune metagenes in the three luminal immune subtypes (low-TIL, high-ISG, and high-TIL) and in non-luminal (HER2-enriched and basal-like) tumors. Each column represents a patient, grouped by immune subtypes; each row represents a gene, grouped by 13 immune pathways. Normalized gene expression value with mean=0 and standard deviation (SD)=1 is indicated by 5 color categories representing the increasing expression level from green to red. LCK: lymphocyte-specific protein tyrosine kinase; Tfh: helper follicular T cell; Tregs: regulatory T cell; NK: natural killer cell; MHC: major histocompatibility complex; STAT1: signal transducer and activator of transcription 1; IF_I: interferon inducible genes; PAM50: green=luminal A, blue=luminal B, grey=basal, black=HER2-enriched.

We also inferred the fractions of 23 immune cell subpopulations in these patients using CIBERSORT. Unlike MCP-counter, CIBERSORT estimates the relative fraction of each cell population in a sample rather than the absolute abundance. Most analyzed immune cell subpopulations had low fractions in our samples. Figure 2b shows the fractions of seven subpopulations with the average fraction >10% across all samples. We found that high-TIL tumors showed significantly higher fractions of CD8+ T cells and tumor-killing M1 macrophages^22^ than those of low-TIL and high-ISG tumors, while they had lower frequencies of tumor-promoting M2 and undifferentiated M0 macrophages (P values see Supplementary Table 5).

**Figure 2:**
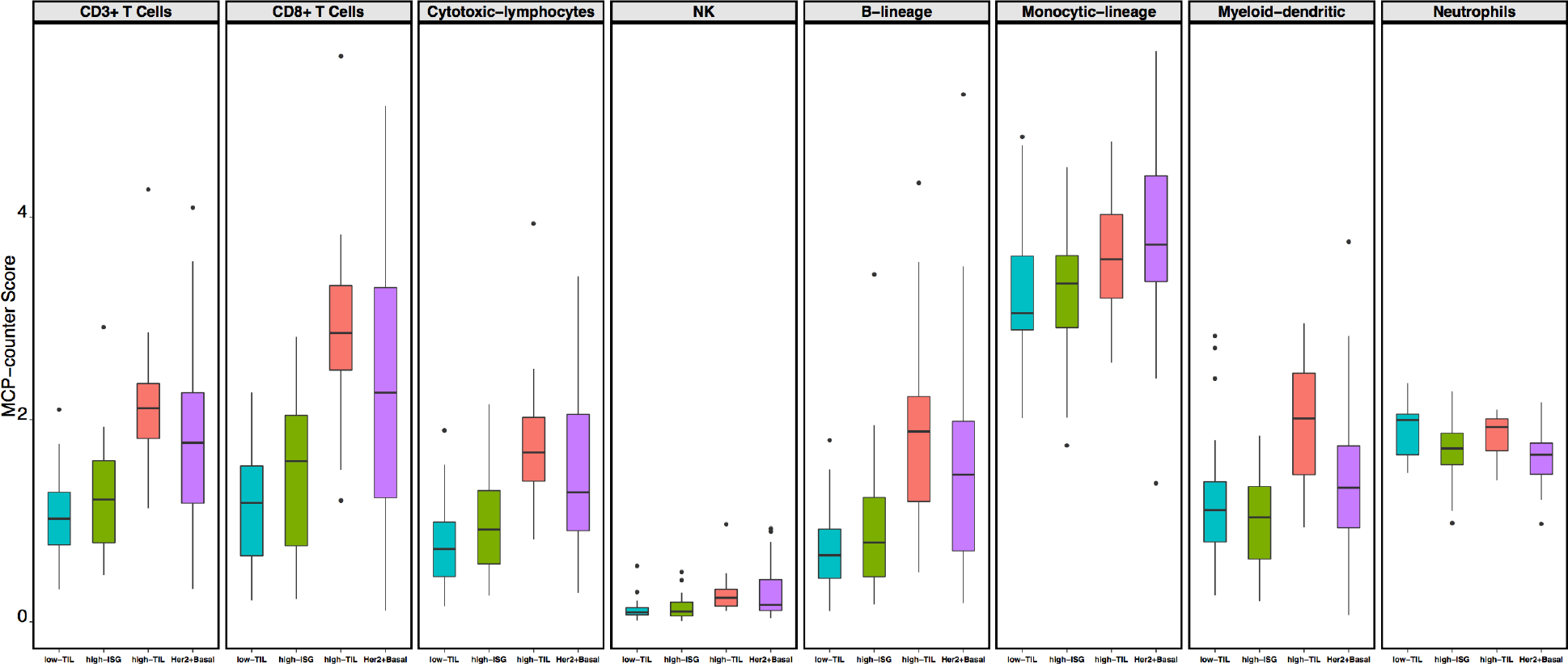
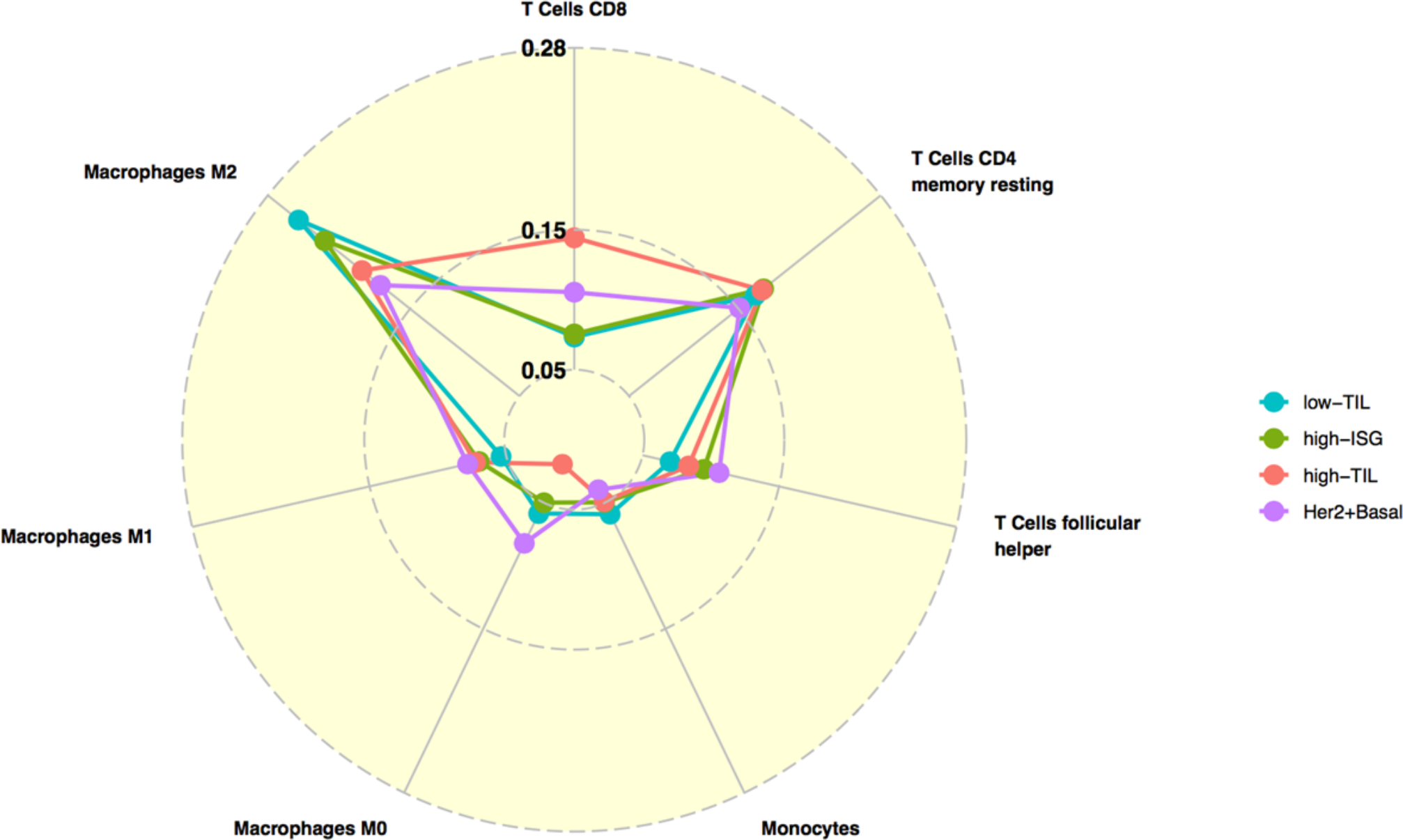
The immune phenotype in the three luminal immune subtypes (low-TIL, high-ISG, and high-TIL) and in non-luminal (HER2-enriched and basal-like) tumors. a) Abundance of eight immune cell subpopulations (estimated by MCP-counter); b) Relative fractions of immune cell populations (inferred by CIBERSORT). Immune cell populations with low fractions (average <10% across all samples) are not shown.

### The presence of luminal immune subtypes was replicated in independent studies

Based on expression levels of the same 130 immune genes used in HKBC, luminal tumors in each TCGA population (Asian, African American, and White) and KBC were similarly assigned to three subtypes using consensus clustering, with the presence of a high-TIL luminal subtype seen in all populations (Figure 3). The pattern was more similar in the three Asian populations, with a more pronounced separation of the high-TIL subtype from the other two subtypes. Consistent with HKBC results, high-TIL tumors in all replication datasets showed higher overall immune score (by ESTIMATE, Figure 3), higher abundance of most immune subpopulations (by MCP-counter, Supplementary Figure 4a), and higher fractions of CD8+ T cells and M1 macrophages (by CIBERSORT, Supplementary Figure 4b). Like HKBC, high-TIL tumors showed upregulation of genes in immune activation and regulation activities (Supplementary Figure 4c), while high-ISG tumors expressed higher levels of ISGs (e.g. *DDX58*) than tumors in the other two luminal immune subtypes (Supplementary Figure 4d).

**Figure 3:**
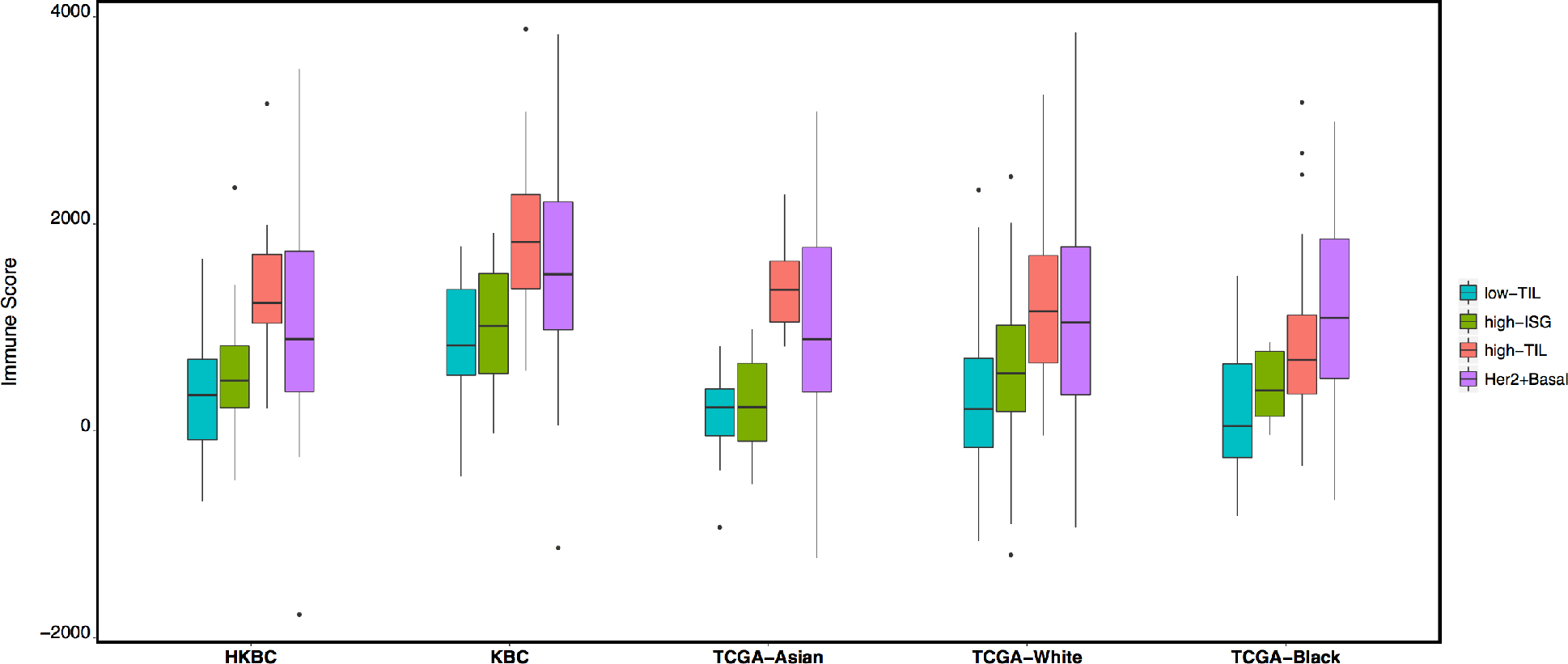
Average immune scores (inferred by ESTIMATE) in the three luminal immune subtypes and non-luminal (HER2-enriched and basal-like) tumors in HKBC, KBC, and TCGA (Asian, African American, and White, separately) datasets.

### Clinical characteristics, breast cancer risk factors, and genomic features associated with immune subtypes

In HKBC, most clinical characteristics or BC risk factors examined, such as tumor grade, nodal status, age at menarche, parity, age at first birth, breastfeeding, and age at menopause, did not vary significantly across immune subtypes (Supplementary Table 1). However, the average body mass index (BMI) was significantly higher in high-TIL (mean = 27.9) than in low-TIL (mean = 24.1) and high-ISG patients (mean = 24.6). The differences remained significant after the adjustment of age, menopausal status, and tumor purity (P=0.0018 for high-TIL vs. low-TIL and P=0.0057 for high-TIL vs. high-ISG). In addition, high-TIL tumors had slightly lower *ESR1* (estrogen receptor alpha) but significantly higher *ESR2* (estrogen receptor beta) expression levels, resulting in a significantly lower *ESR1*/*ESR2* ratio (P=0.001) compared with low-TIL and high-ISG tumors (Figure 4a). The association between the low *ESR1/ESR2* ratio and the high-TIL subtype was consistently seen in all TCGA populations (Supplementary Figure 5a).

High-TIL patients tended to be younger than patients with low-TIL tumors in HKBC as well as in the replication datasets (Supplementary Figure 5b), although the difference was significant only among the TCGA Whites (P=0.018). The short follow-up time in HKBC prohibited us from assessing the prognostic outcome in relation to the immune subtypes. We therefore conducted survival analysis using TCGA data of 905 BC patients. We combined all ethnicity groups because few deaths occurred among Asian or African American patients. As shown in Supplementary Figure 5c, the high-TIL subtype was associated with the best 10-year overall survival among all subtypes (P=0.008), although the difference became non-significant after the adjustment for age at diagnosis and stage (hazard ratio [HR]=0.6, 95% confidence interval [CI]=0.26-1.4, P=0.22). The attenuation of the significance was likely due to younger ages in the high-TIL subtype as stage did not differ significantly across luminal immune subtypes (P=0.72).

To evaluate the possible contribution of germline variation in *APOBEC3B* to immune profiles and mutational events, we genotyped a SNP (rs12628403) that is a proxy for the *APOBEC3B* deletion in germline DNA^23^. In HKBC, the frequency of the rs12628403-C allele that tags the 30kb deletion (44.7% among 76 luminal patients and 40.4% among all 114 patients with genotyping data) was similar to what was reported in East Asian populations^18^. We found the expected associations between the *APOBEC3B* deletion and decreased levels of *APOBEC3B* expression in both tumor and normal tissue, validating SNP rs12628403 as a proxy for *APOBEC3B* deletion (Supplementary Figure 6). Although high-TIL patients had a slightly higher frequency of the deletion allele than the other luminal immune subtypes, the difference was not significant (P=0.93, Figure 4b). In addition, the expression level of *APOBEC3A_B*, which is a hybrid transcript resulting from the *APOBEC3B* deletion, did not vary significantly by luminal immune subtypes (P=0.36). Further, the ESTIMATE immune scores did not vary across different genotypes of SNP rs12628403(P=0.56). Similar results were obtained in the analysis based on all tumor subtypes. In TCGA Whites, the homozygous deletion of *APOBEC3B* was very rare; only 2 of 329 patients with genotyping data were homozygote and neither of them was in the high-TIL subtype (Figure 4b).

In an exploratory analysis of a subset of luminal tumors with both RNA-Seq and WES data (n=59), we found that, after age adjustment, high-TIL tumors were associated with a higher nonsynonymous mutation burden (P=0.014, compared to low-TIL tumors, Figure 4c) and a higher frequency of *APOBEC*-signature mutations (mean 23.6%) compared with low-TIL (7.6%, P=0.04) and high-ISG (8.3%, P=0.05) tumors. Notably, all *TP53* mutations (n=8, Figure 4d) observed among luminal patients occurred in high-ISG tumors. The similar enrichment of *TP53* mutations in high-ISG tumors was also seen in TCGA Whites (P=0.006, Figure 4d). The frequency of *PIK3CA* mutations did not vary significantly by immune subtypes in HKBC but showed a slight increase in high-TIL tumors in TCGA Whites (P=0.031 compared to low-TIL tumors).

**Figure 4:**
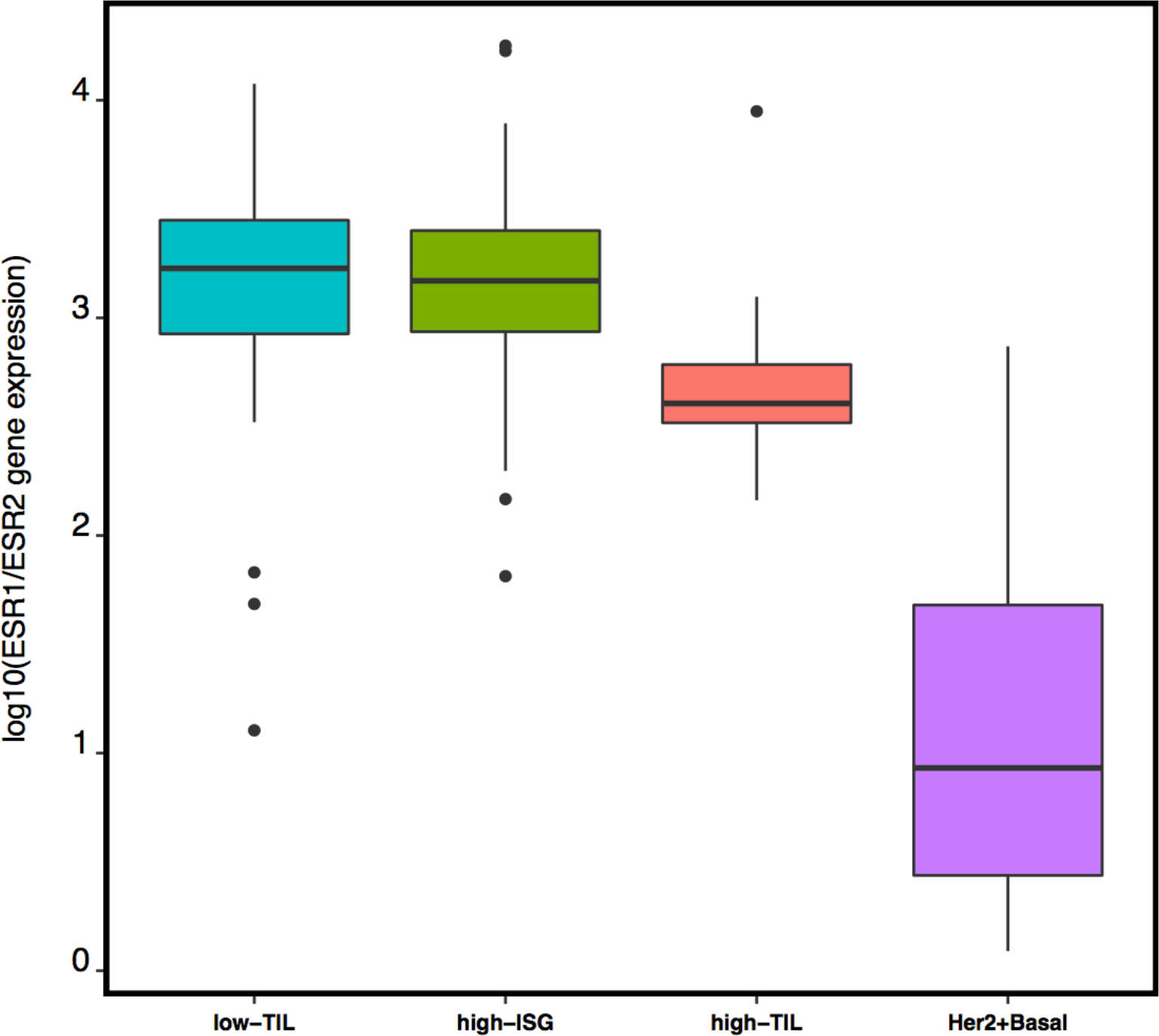
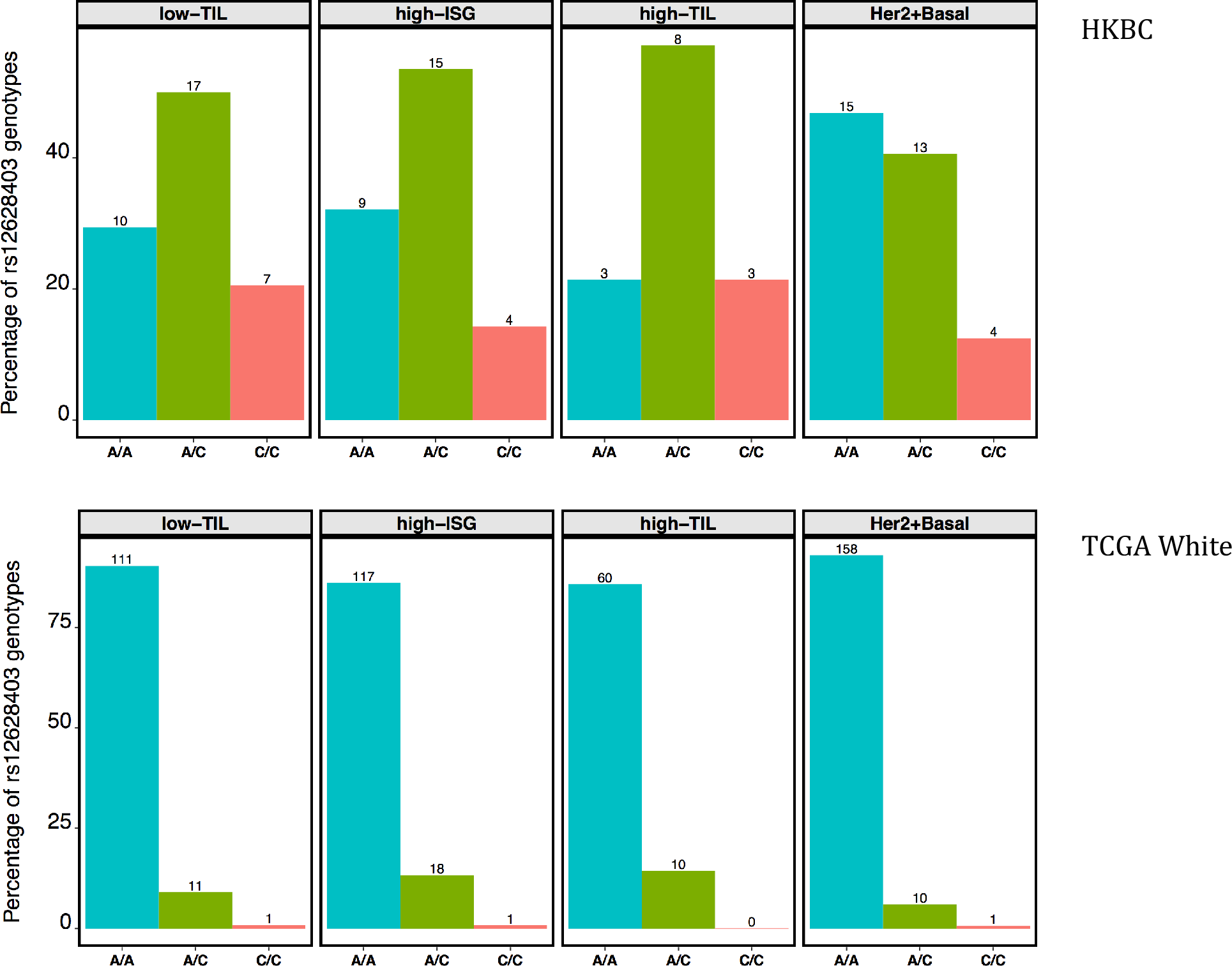
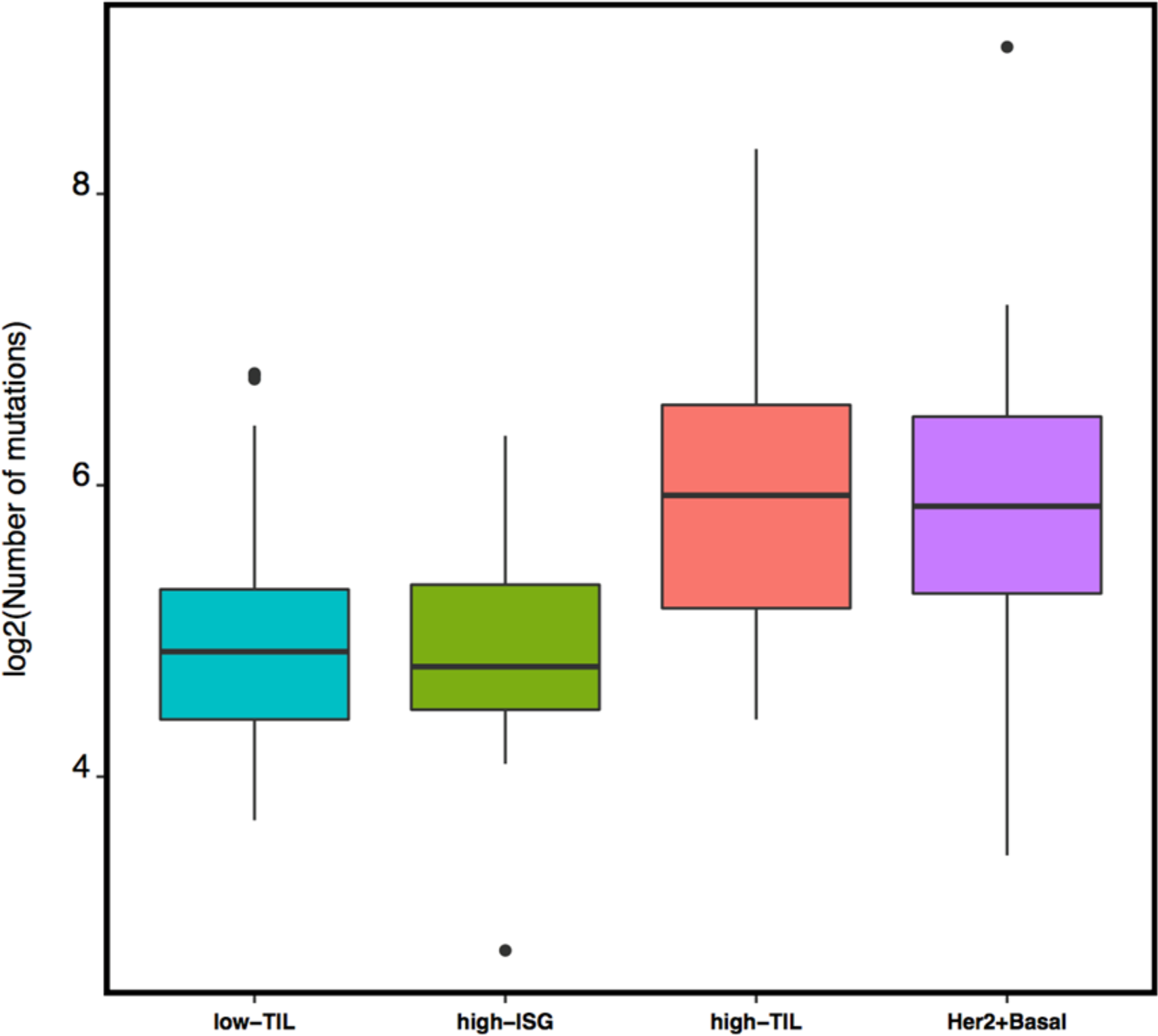
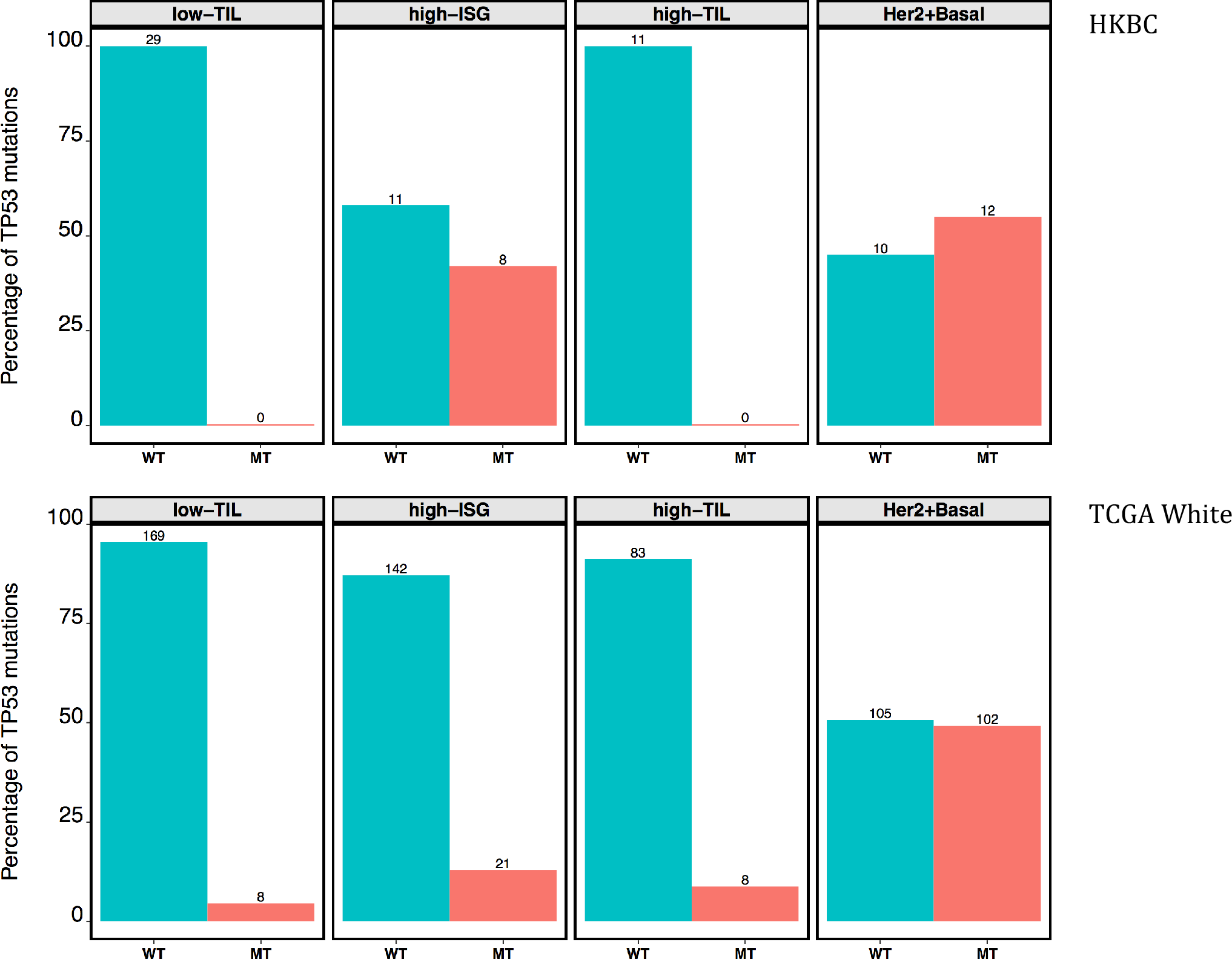
Genomic features associated with different immune subgroups. a) *ESR1* and *ESR2* expression ratio (log scale); b) Number of patients with the germline *APOBEC3B* deletion tagged by rs12628403-C allele in HKBC and TCGA Whites (number of patients in each genotype category indicated above each bar); c) Nonsynonymous mutation burden (log scale); d) Frequency of nonsynonymous *TP53* mutations in HKBC and TCGA White patients (number of patients in each mutation group indicated above each bar).

### Comparison to matched normal tissue suggested T cell activation in high-TIL tumors only

In our HKBC data neither abundance nor fractions of the examined immune cell populations in paired normal breast tissue varied significantly across the three luminal immune subtypes (Supplementary Figure 7), suggesting that the distinguishing TIL levels between high-TIL and other tumors were not driven by the differences in their systematic normal TIL levels. We compared the MCP abundance of the eight immune cell populations in tumor (T) and matched normal (N) tissues for each immune subtype. Low-TIL and high-ISG overall showed similar patterns of immune discrepancy between T and N, while patterns of high-TIL were more similar to those of non-luminal patients (Figure 5). Specifically, while low-TIL and high-ISG tumors showed either no change or lower abundance of immune cell populations (such as cytotoxic lymphocytes) compared to their paired normal tissue, high-TIL, like non-luminal tumors, had significantly higher abundance scores of CD3+ T cells, CD8+ T cells, and B lineage cells compared with the paired normal tissue (T - N difference > 0, Figure 5; P-value of CD8+ T cells = 0.0002 and 0.0253 for high-TIL and non-luminal patients; other P values see Supplementary Table 6). These observations indicate a tumor-derived activation of specific immune responses in high-TIL and non-luminal tumors but not in other luminal tumors.

**Figure 5:**
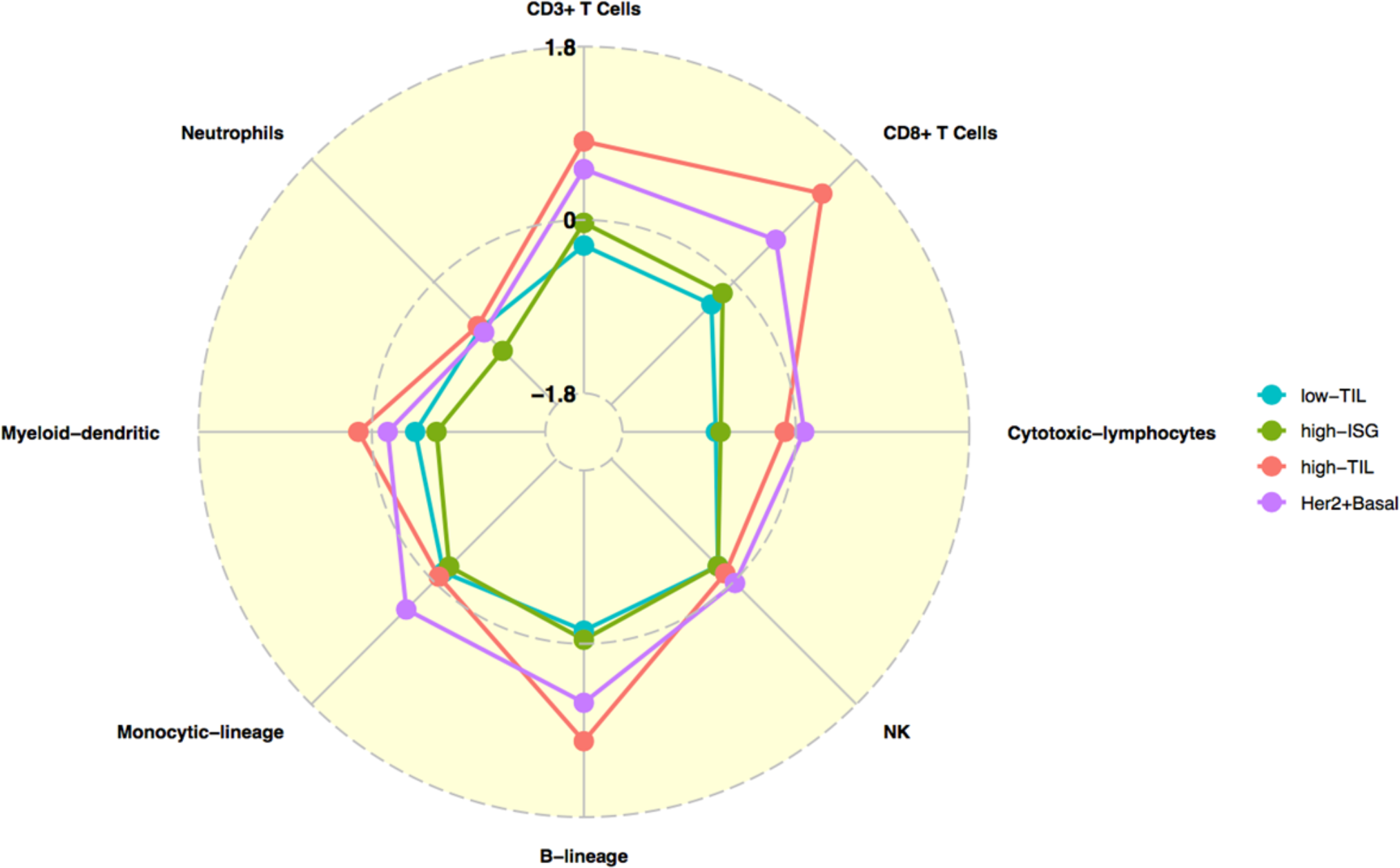
Comparison of the abundance of eight immune cell subpopulations (estimated by MCP-counter) between paired tumor and normal tissue (N=80) for the three luminal immune subtypes and non-luminal (HER2-enriched and basal-like) patients in HKBC, respectively. Each dot represents the mean difference between each tumor and normal pair (T-N); 0: no difference, >0: higher in tumor than normal tissue; <0: lower in tumor than in normal tissue.

## Discussion

In this study of immune and genomic characterization of luminal breast cancer patients, we identified three immune subtypes of luminal breast tumors displaying distinct patterns of immune gene expression with associated genomic features. One luminal subtype (high-TIL) exhibited an activated immune phenotype that is comparable to that of non-luminal (HER2-enriched and basal-like) tumors. These high-TIL tumors were predominantly luminal-A and were found in approximately one-sixth of luminal patients in the HKBC study. Like non-luminal tumors, high-TIL luminal tumors had higher expression of immune checkpoint genes, higher abundance and fractions of CD8+ lymphocytes, and higher ratio of M1/M2 macrophages. They also shared genomic features of non-luminal tumors, such as a higher mutation burden, higher fractions of *APOBEC*-signature mutations, and lower *ESR1/ESR2* expression ratio. High-TIL luminal patients also appear to have a better prognosis and higher BMI compared to other luminal patients in our study. In addition to high-TIL, we also identified a luminal subtype (high-ISG) that was characterized by increased expression of ISGs and enrichment for *TP53* mutations. These immune subtypes were replicated in independent datasets. Our findings suggest that immune gene expression and associated genomic features may reveal additional heterogeneity in luminal breast cancer patients beyond the current luminal A/B classification.

Our immune subtyping of TCGA breast cancer patients showed modest correlation with the six immune subtypes identified by Thorsson *et al.* in a TCGA Pan-Cancer analysis^13^. Specifically, our low-TIL, high-ISG, and high-TIL subtypes were enriched with C1 (wound healing), C2 (IFN-γ dominant), and C3 (inflammatory) subtypes, respectively, in the Pan-Cancer study. The modest correlation is not surprising since the clustering analysis is heavily influenced by the input genes and subjects. In contrast to the large number of immune genes (~3,000) and a heterogeneous mixture of cancer types used in the Pan-Cancer analysis, our clustering analysis included a more focused immune gene panel (130 genes) and was restricted to luminal breast tumors, which may better capture the variation in tumor immunogenicity within luminal breast tumors. Indeed, the concordance index (C-index), which evaluates the concordance of the actual and predicted survival outcomes for TCGA luminal breast cancer patients, was slightly higher for our luminal immune subtypes (0.60) than the one for the Pan-Cancer immune subtypes (0.56).

Previous studies suggested that the high expression of an alternative ER isoform, *ESR2* (encoding ERβ), was associated with favorable BC prognosis and that the association might depend on the ratio of *ESR1* and *ESR2* (ERα and ERβ)^24,25^. Consistently, we observed that patients with increasing *ESR1/ESR2* ratio tended to have poorer survival (HR=1.5, 95% CI=0.7-3.3, P=0.27, adjusting for age and stage) in TCGA luminal patients. Interestingly, in the current study, we found that high-TIL tumors had a significantly lower *ESR1/ESR2* ratio as compared to low-TIL and high-ISG tumors in both HKBC and replication datasets. Our findings suggest that *ESR* expression, particularly *ESR2* expression, may relate to immune gene regulations in luminal breast tumors and this association may explain the previously reported favorable prognosis associated with ERβ expression.

We also identified a unique subtype (high-ISG) of luminal tumors that had lower scores in most immune pathways but showed higher expression of *STAT1* and other ISGs (such as *DDX58*) even as compared to high-TIL tumors. Unlike low-TIL and high-TIL subtypes that comprised predominantly luminal-A tumors, high-ISG patients were enriched with luminal-B tumors.

Interestingly, all *TP53* mutations among luminal patients occurred in the high-ISG subtype in HKBC. Previous studies demonstrated that *TP53* mutations were associated with an immune activated phenotype when all molecular subtypes were analyzed together, which is expected since *TP53* mutations are more prevalent in non-luminal than in luminal tumors. Our data suggest that *TP53* mutations may be specifically related to the activation of IFN-signaling. p53 has been reported to inhibit STAT1, a key transcription factor in the JAK-STAT pathway that drives the expression of ISGs and pro-inflammatory cytokines^26^. It is therefore possible that the deficiency of p53 caused by *TP53* mutations may contribute to the overexpression of *STAT1* and ISGs. Alternatively, *TP53* mutations and overexpression of ISGs may be the combined result of analogous DNA damaging events since induction of ISGs may occur even without viral infection in response to DNA damage^27^. The association between *TP53* mutations and high-ISG was also seen in TCGA EA. Our results suggest that the relationship between immune composition and genomic determinants might be more complex than we previously appreciated.

In our study, we did not find a significant association between the germline *APOBEC3B* deletion and luminal immune subtypes. Similarly, the immune scores did not vary significantly by the deletion genotype, either in luminal or in all patients. The previously observed association between the deletion and immune activation was based on data from TCGA and METABRIC, in which the frequency of the homozygous deletion was very low^19,20^ and the results were driven by comparing the heterozygotes to the wild type. Although our evaluation was limited by the overall small sample size, the higher frequency of the deletion in this Asian population allowed us to examine both heterozygous and homozygous genotypes. Results based on our study do not support the hypothesis that the germline *APOBEC3B* deletion polymorphism is the driving force for immune activation in breast tumors^19,20^.

Taking advantage of our rich collection of epidemiologic data in HKBC, we examined several established breast cancer risk factors in relation to the immune subtypes and found a significant association between higher BMI and the high-TIL luminal subtype. The average BMI was more than 3 units higher in high-TIL patients compared with other luminal patients and the differences remained significant after the adjustment for potential confounders such as age at diagnosis, menopausal status, and tumor purity. The association was not seen in KBC, likely due to the lack of BMI variation in such a young cohort. Consistent with our finding in HKBC, a recent study reported a significant association between higher expression of CD8+ T-cell signatures and increasing BMI in 1,154 breast cancer patients from the Nurses’ Health Study^28^. The link between obesity and breast cancer involves multiple mechanisms that may interplay with each other such as chronic inflammation, estrogen production, growth factor stimulation, and altered metabolism^29^. Future large studies are warranted to follow up this observation.

In contrast to tumors, immune gene expression in adjacent normal tissue did not vary significantly across the three luminal immune subtypes, suggesting that high-TIL patients did not have high immune activation. Using gene expression data based on BC patients in Norway, Quigley et al. previously showed that cytotoxic lymphocyte (CTL) pathway scores were higher in tumors than in matched adjacent normal tissue, particularly for ER-negative tumors^30^. We found that high-TIL patients also showed significantly higher levels of CD3+ T cells, CD8+ T cells, and B lineage cells in their tumors compared with normal tissues. These findings suggest that tumor-intrinsic events might drive the immune activation in a similar manner in ER-negative and high-TIL luminal tumors. In fact, consistent with what was reported by several previous studies^31,32^, we found that higher burden of nonsynonymous mutations and *APOBEC*-signature mutations might act as potential contributors to the increased immune response.

The strengths of our study include a comprehensive collection of clinical and exposure information and a detailed evaluation of immune composition for both tumors and paired normal tissue in an Asian population, and the replication of findings in independent datasets. The major limitation is the small sample size, which limited the power to identify genomic determinants of distinct immune phenotypes. Additionally, since we collected frozen breast tissue from recently diagnosed patients, the follow-up time is insufficient to evaluate the associations between the immune subtypes with prognostic outcomes. Large TIL studies of luminal BC with treatment and outcome data are warranted to follow up on our findings. In summary, we identified three immune subtypes of luminal breast tumors displaying distinct patterns of immune gene expression with associated genomic features. If confirmed, these findings may have important clinical implications in improving luminal BC stratification for precision oncology treatment^1,5,10–12^.

## Methods

### Participants and Samples

We analyzed data and biospecimens collected from a hospital-based breast cancer case-control study in Hong Kong as previously described^33^. In brief, fresh frozen breast tumors and paired normal tissues were collected from newly diagnosed breast cancer patients in two HK hospitals between 2013 and 2016. Patients with pre-surgery treatment were excluded from the study.

Clinical characteristics and breast cancer risk factors were obtained from medical records and questionnaire. Paired tumor and histologically normal breast tissue samples were processed for pathology review at the Biospecimen Core Resource (BCR), Nationwide Children's Hospital, using modified TCGA criteria^34^. Tumors with >50% tumor cells and normal tissues with 0% tumor cells were included for DNA/RNA extraction. The study protocol was approved by ethics committees of the Joint Chinese University of Hong Kong-New Territories East Cluster, the Kowloon West Cluster, and the National Cancer Institute. Written informed consent was obtained prior to the surgery for all participants.

### Transcriptome sequencing, PAM50 classification, and immune composition

RNA sequencing (RNA-Seq) data was generated in 139 tumors and 92 histologically normal breast tissue paired samples that passed standard QC metrics at Macrogen Corporation on Illumina HiSeq4000 using TruSeq stranded RNA kit with Ribo-Zero for rRNA depletion and 100-bp paired-end method. Gene expression was quantified as TPM (transcript per million) using RSEM^35^ and log_2_TPM was used for statistical analyses. PAM50 subtype was defined by an absolute intrinsic subtyping (AIMS) method^36^. A comprehensive characterization of immune cell composition in both tumor and paired normal breast tissue was achieved by using three computational algorithms: ESTIMATE^37^, CIBERSORT^38^, and MCP-counter^39^. While ESTIMATE (for overall infiltration of immune cells) and MCP-counter (for eight immune cell subpopulations) both measure the abundance of immune cells in a given sample, CIBERSORT estimates intra-sample proportions of 23 immune cell subpopulations.

### Whole-exome sequencing (WES) and mutation analyses

WES was performed on 104 paired tumor and normal samples (40% from blood or saliva and 60% from histologically normal breast tissue) at the Cancer Genomics Research Laboratory (CGR), NCI, using SeqCAP EZ Human Exome Library v3.0 (Roche NimbleGen, Madison, WI) for exome sequence capture. The captured DNA was then subject to paired-end sequencing utilizing Illumina HiSeq2000. 59 of them also had RNA-Seq data. The average sequencing depth was 106.2× for tumors and 47.6× for the paired normal tissues. Somatic mutations were called using four callers and the analyses were based on mutations called by 3 or more of 4 established callers (MuTect^40^, MuTect2 (**GATK tool**), Strelka^41^, and TNScope by Sentieon^42^). Mutation signatures were estimated using the previously published method^43^. SNP rs12628403, which is a proxy for the *APOBEC3B* deletion (r^2^=1.00 in Chinese from Beijing (CHB) in HapMap samples), was genotyped in germline DNA with a custom TaqMan assay as previously described^23^.

### Replication datasets

We analyzed two available, independent datasets to replicate our findings: 564 luminal patients in TCGA^3^and 112 luminal patients in a Korean BC genomic study (KBC)^44^. We analyzed TCGA Asians (n=29, mean age: 51 years), African Americans (AA, n=72, mean age: 58 years), and European ancestry (EA, n=463, mean age: 60 years) separately. PAM50 was called using the same AIMS method for each TCGA sample as it was used in HKBC. KBC patients were much younger, with a mean age at diagnosis of 40 years. PAM50 subtype and mutation calling for KBC were previously detailed^44^. Immune classification and composition across all datasets (HKBC, TCGA, and KBC) were analyzed using the same methods.

### Statistical Analysis

The consensus clustering was conducted using ConsensusClusterPlus^45^. The ANOVA test was used to compare mean differences across the luminal immune subtypes for immune cell populations and their immune scores. Logistic regression was used to assess the associations between the immune subtypes (outcome) and transcriptomic features, genomic alterations, patient characteristics, and breast cancer risk factors, with the adjustment for age at diagnosis. The Kaplan–Meier method was used to assess overall survival among patients, stratified by immune subtypes. A multivariable Cox proportional hazards model was also used to test the differences in survival across immune subtypes with the adjustment of age at diagnosis and tumor stage. Since most of our analyses were exploratory, we did not adjust for multiple testing. All statistical tests were two-sided and performed using SAS version 9.3 (SAS Institute, Cary, NC, USA) or R version 3.4.4 (R Foundation for Statistical Computing, Vienna, Austria).

## Author Contributions

Dr. Yang had full access to the data in the study and takes full responsibility for the integrity of the data and the accuracy of the data analysis.

*Study concept and design:* Zhu, Tse, Garcia-Closas, Chanock, Yang.

*Acquisition, analysis, or interpretation of data:* Zhu, Tse, Wang, Koka, Zhang, Abubakar, Zhang, Tang, Kan, Lee, Park, Nam, Wang, Sun, Prokunina-Olsson, Shi, Yang.

*Drafting of the manuscript:* Zhu, Yang.

*Critical revision of the manuscript for important intellectual content:* All authors.

*Statistical analysis:* Zhu, Wang, Koka, Zhang, Zhang.

*Obtained funding:* Tse, Yang.

*Administrative, technical, or material support:* Lee, Wang, Wu, Tsang, Cha, Law, Li, Li, Wu, Liu, Huang, Wang, Jones, Zhu, Hutchinson, Hicks.

*Study supervision:* Yang.

## Conflict of Interest Disclosures

None.

## Acknowledgements

This research was supported by the Intramural Research Program of the National Institutes of Health, National Cancer Institute, Division of Cancer Epidemiology and Genetics, and Research Grants Council (grant number 474811 to Dr. Tse), Hong Kong SAR.

