## Supplementary figures for "Immune gene expression profiling reveals heterogeneity in luminal breast tumors"

**Supplementary Figure 1** - Expression levels of immune-related genes in the three luminal immune subtypes and non-luminal (HER2-enriched and basal-like) tumors in HKBC.

**Supplementary Figure 2** - MCP-counter scores of eight immune cell subpopulations in the three luminal immune subtypes and non-luminal (HER2-enriched and basal-like) tumors with the adjustment of tumor purity in HKBC.

**Supplementary Figure 3** - Representative images of H&E stained slides showing low-TIL (A) and high-TIL (B) luminal tumors; red arrow: tumor area; yellow arrow: area with immune infiltration.

**Supplementary Figure 4** - Replication of luminal immune subtypes in TCGA and KBC datasets. a) MCP-counter scores for eight immune cell subpopulations; b) Relative fractions of immune cell subpopulations by CIBERSORT (cell populations with extremely low fractions were not shown); c) High-TIL tumors showed upregulation of genes in immune activation and regulation activities than tumors in the other two luminal immune subtypes; d) High-ISG tumors expressed higher levels of ISG genes than tumors in the other two luminal immune subtypes.

**Supplementary Figure 5** - Replication of genomic features associated with luminal immune subtypes in TCGA and KBC datasets. a) *ESR1*/*ESR2* ratios; b) age at diagnosis; c) 10-year overall survival.

**Supplementary Figure 6** - Expression of *APOBEC3B* in normal and tumor tissue in relation to the polymorphic germline *APOBEC3B* deletion represented by rs12628403-C allele in HKBC.

**Supplementary Figure 7** - MCP-counter scores of eight immune cell subpopulations in adjacent normal breast tissue in the three luminal immune subgroups and non-luminal (HER2-enriched and basal-like) patients of HKBC.

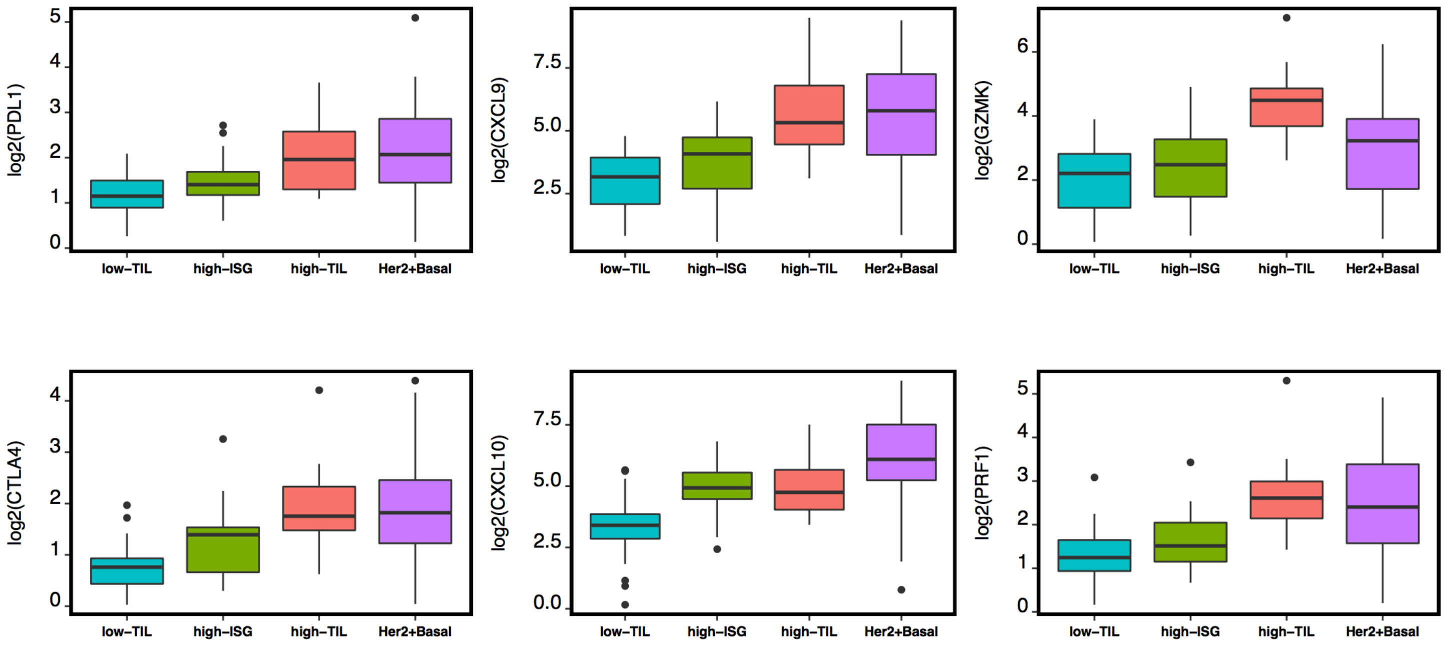


**Supplementary Figure 1**

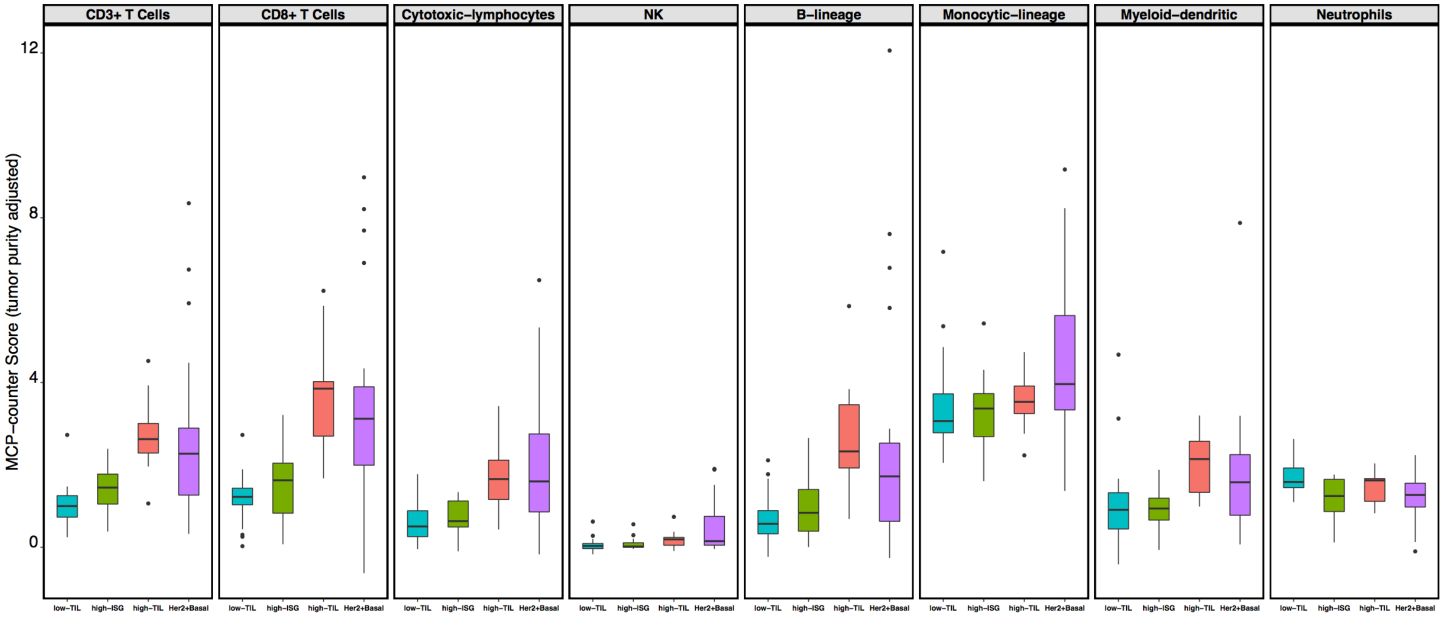


**Supplementary Figure 2**

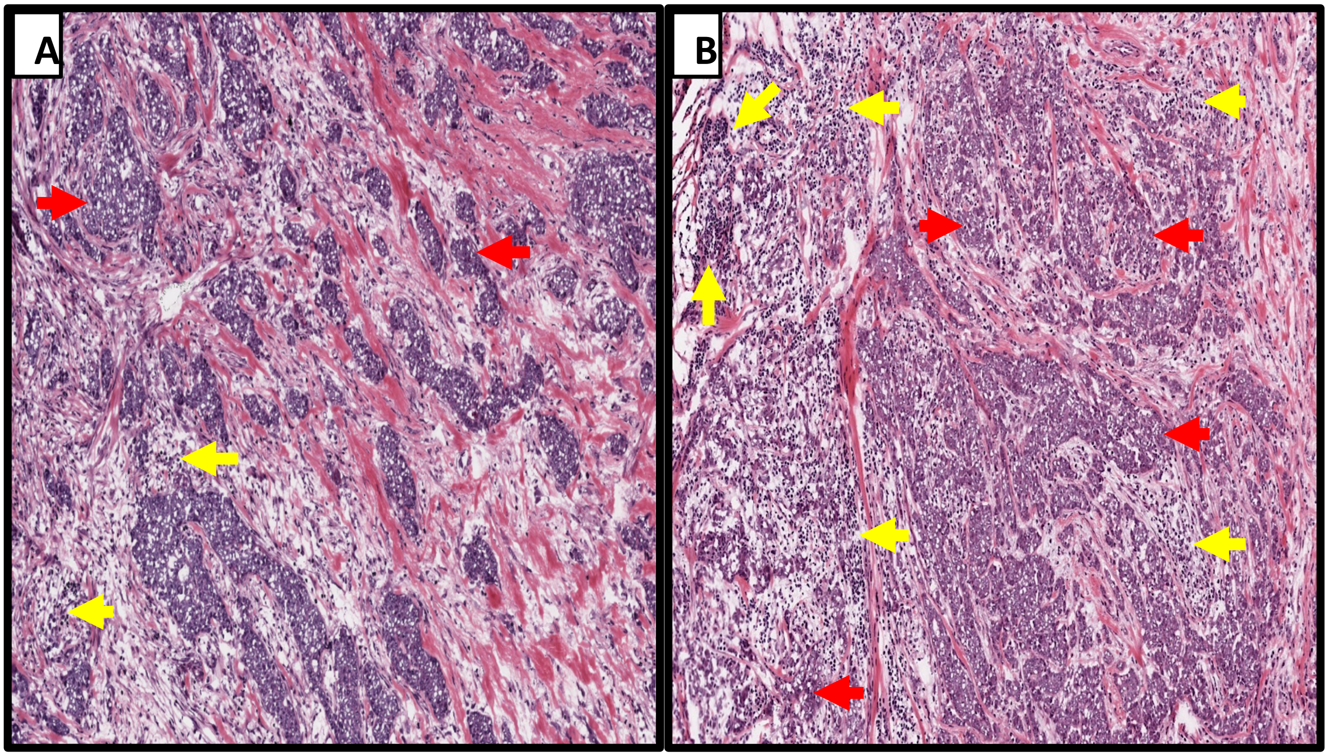


**Supplementary Figure 3**

### TCGA Asian

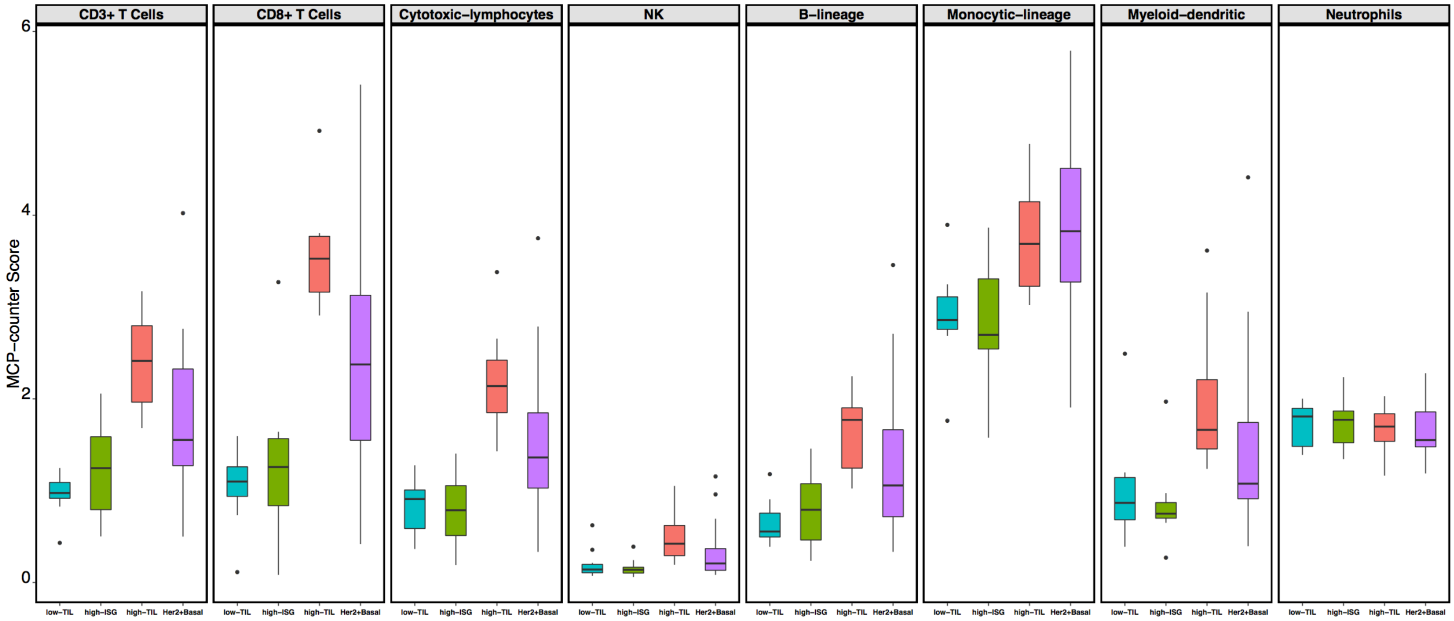


### TCGA White

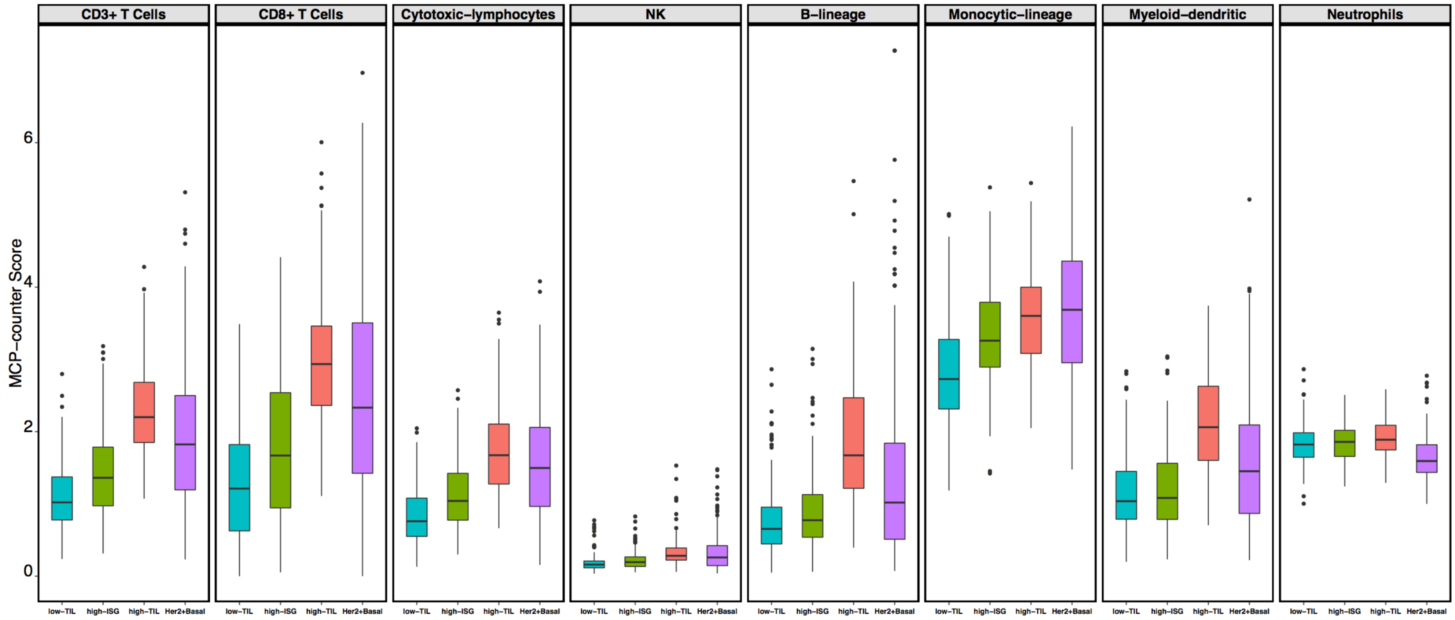


**Supplementary Figure 4.a (*to be continued*)**

### TCGA Black

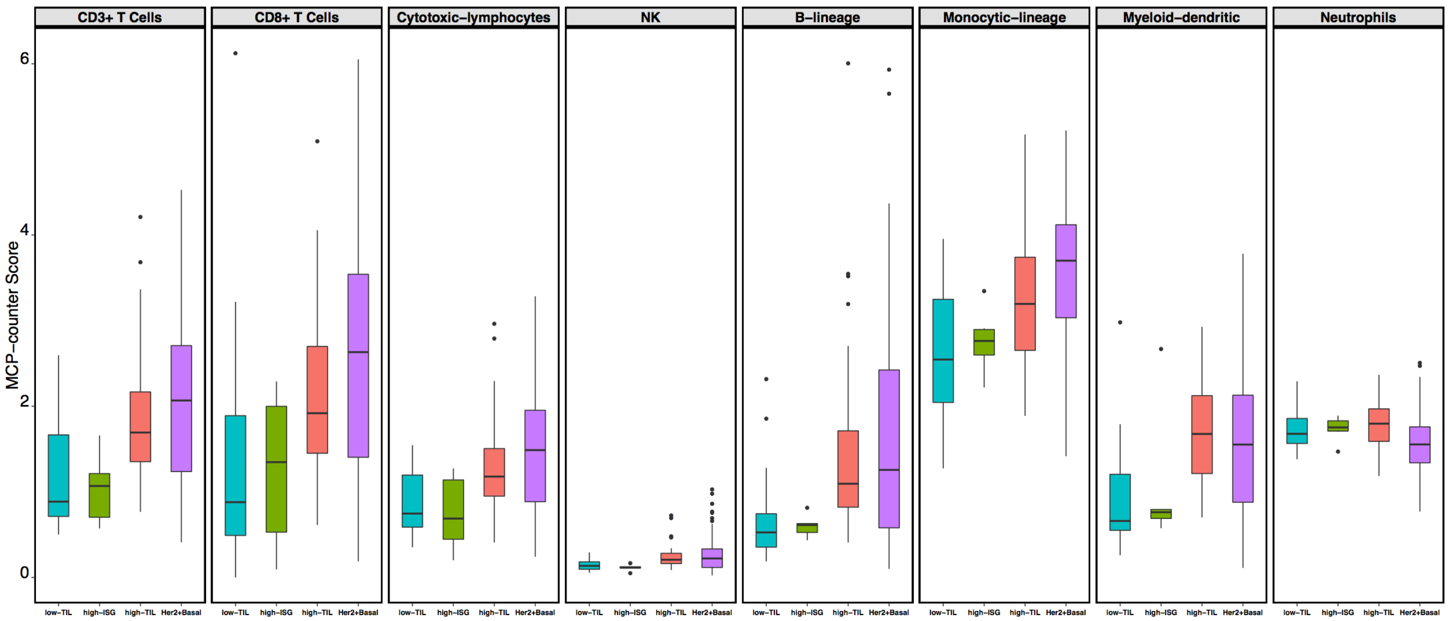


### KBC

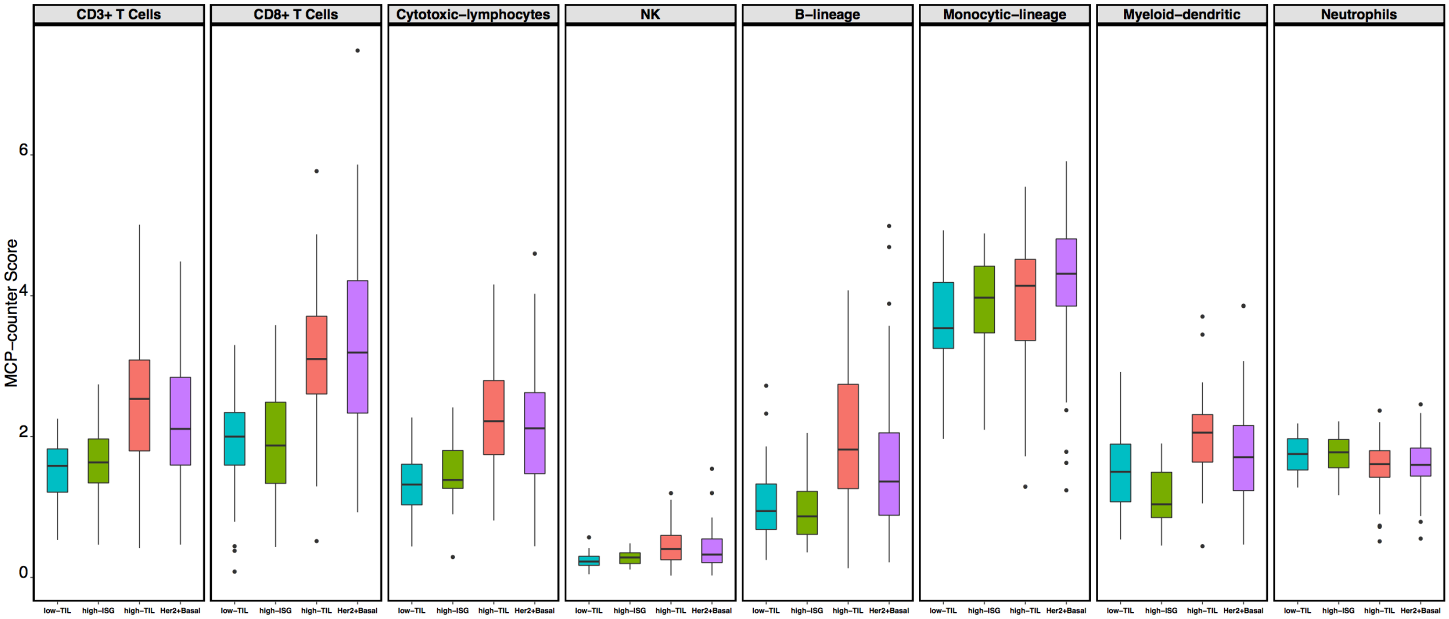


**Supplementary Figure 4.a**

### TCGA Asian

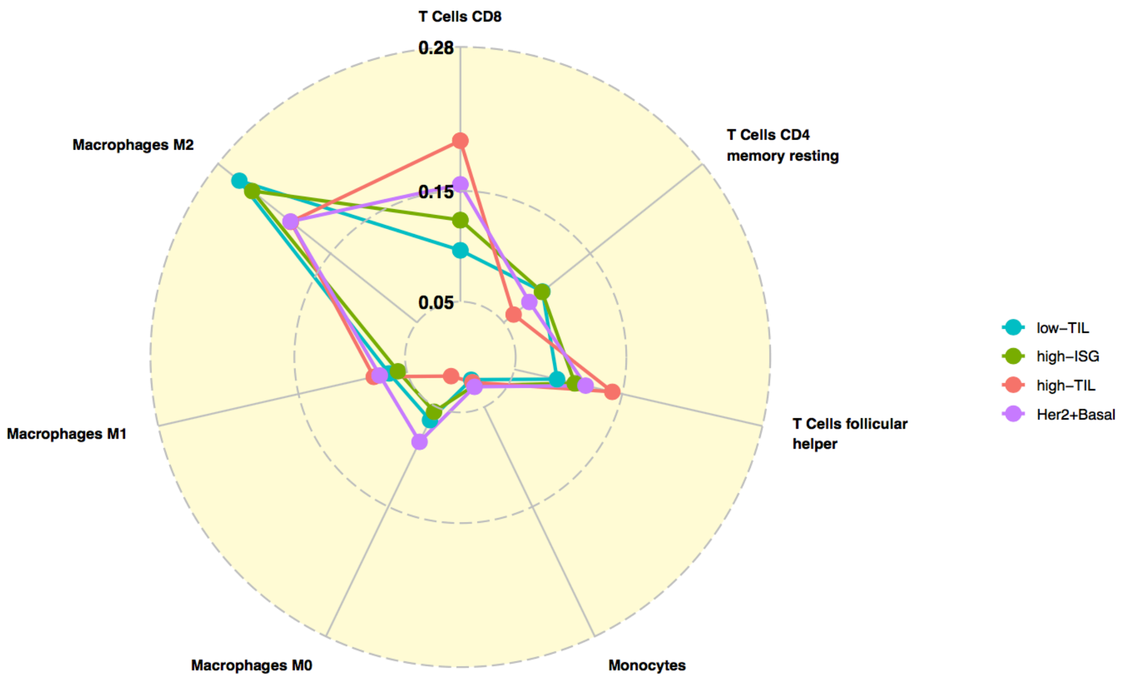


### TCGA White

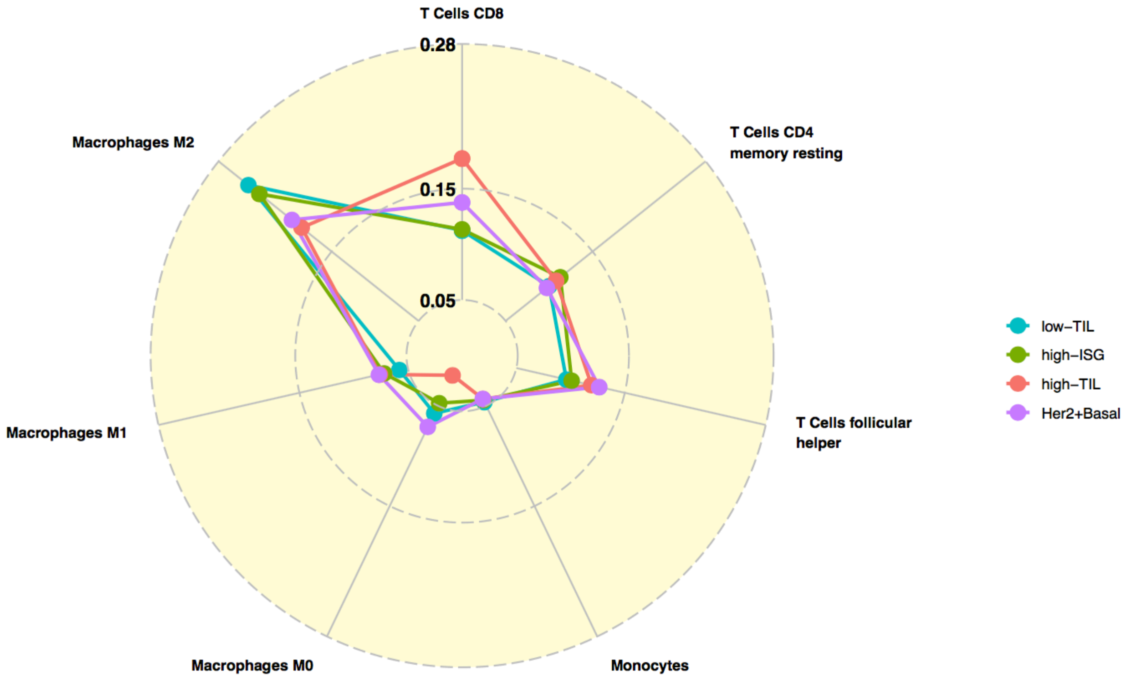


**Supplementary Figure 4.b (*to be continued*)**

### TCGA Black

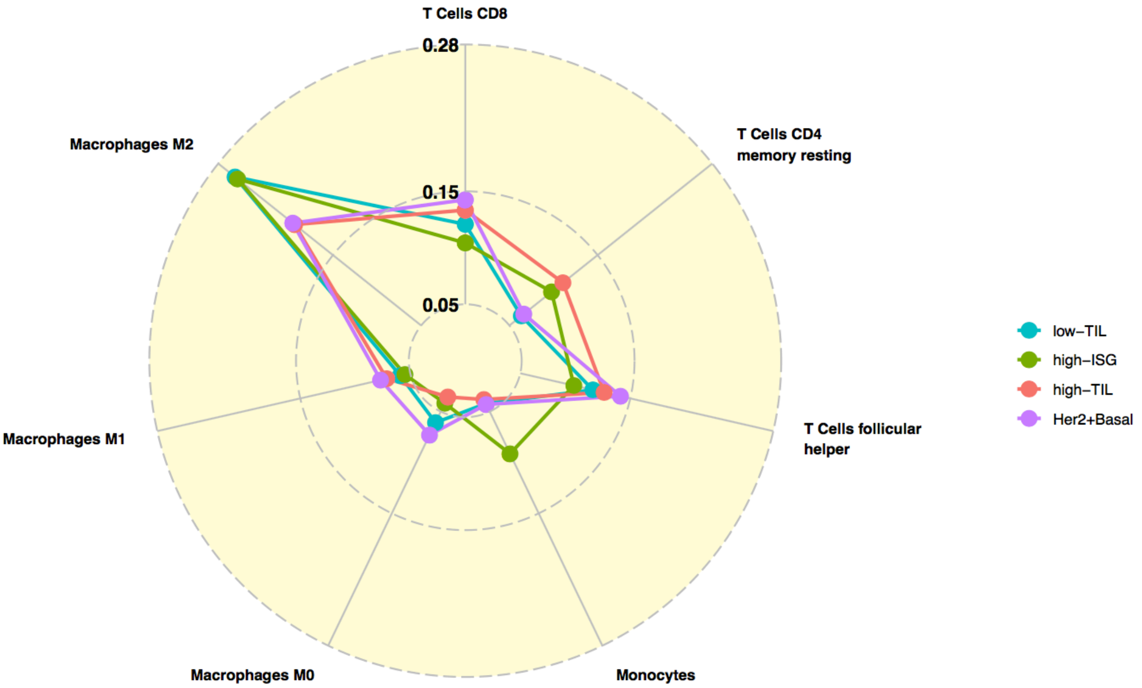


### KBC

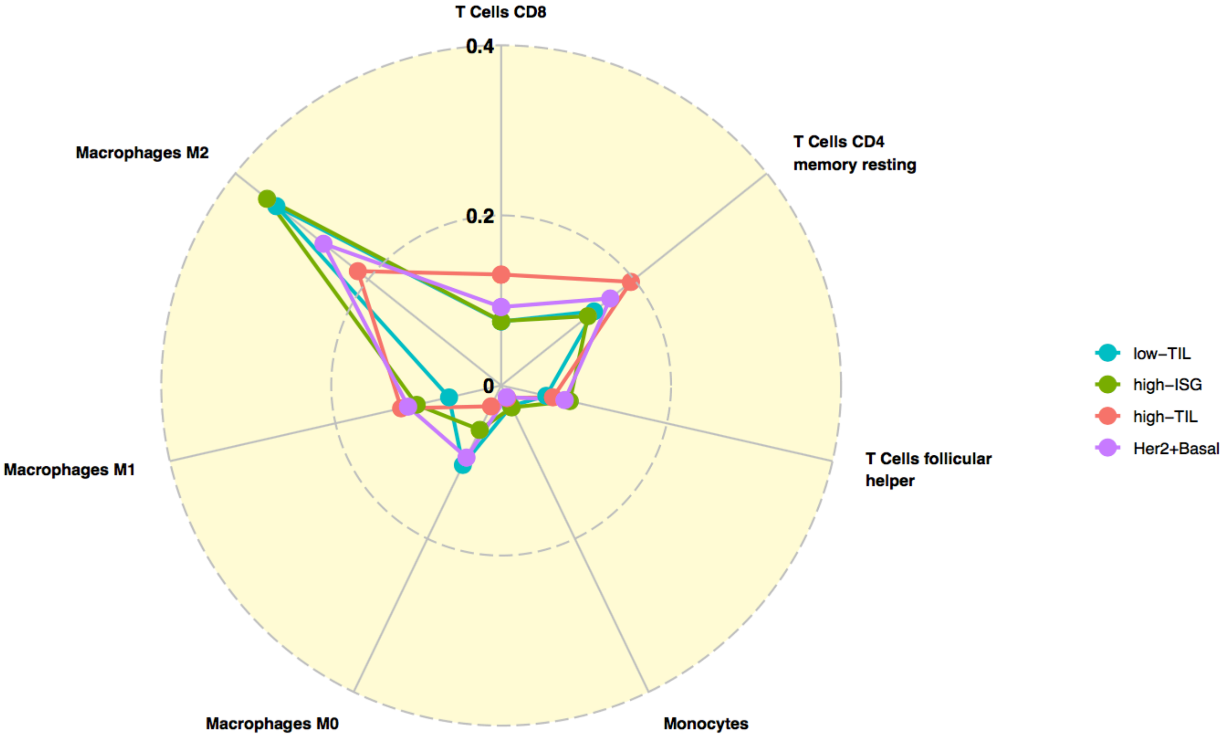


**Supplementary Figure 4.b**

### TCGA

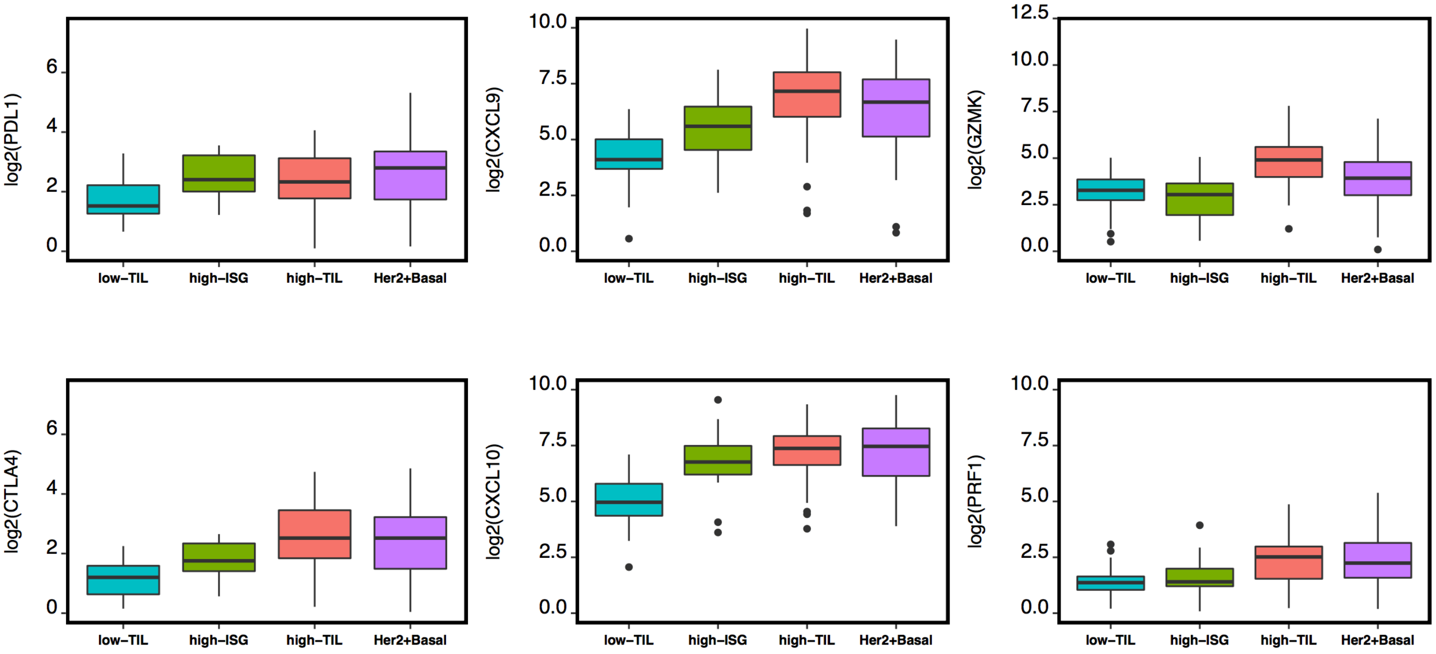


### KBC

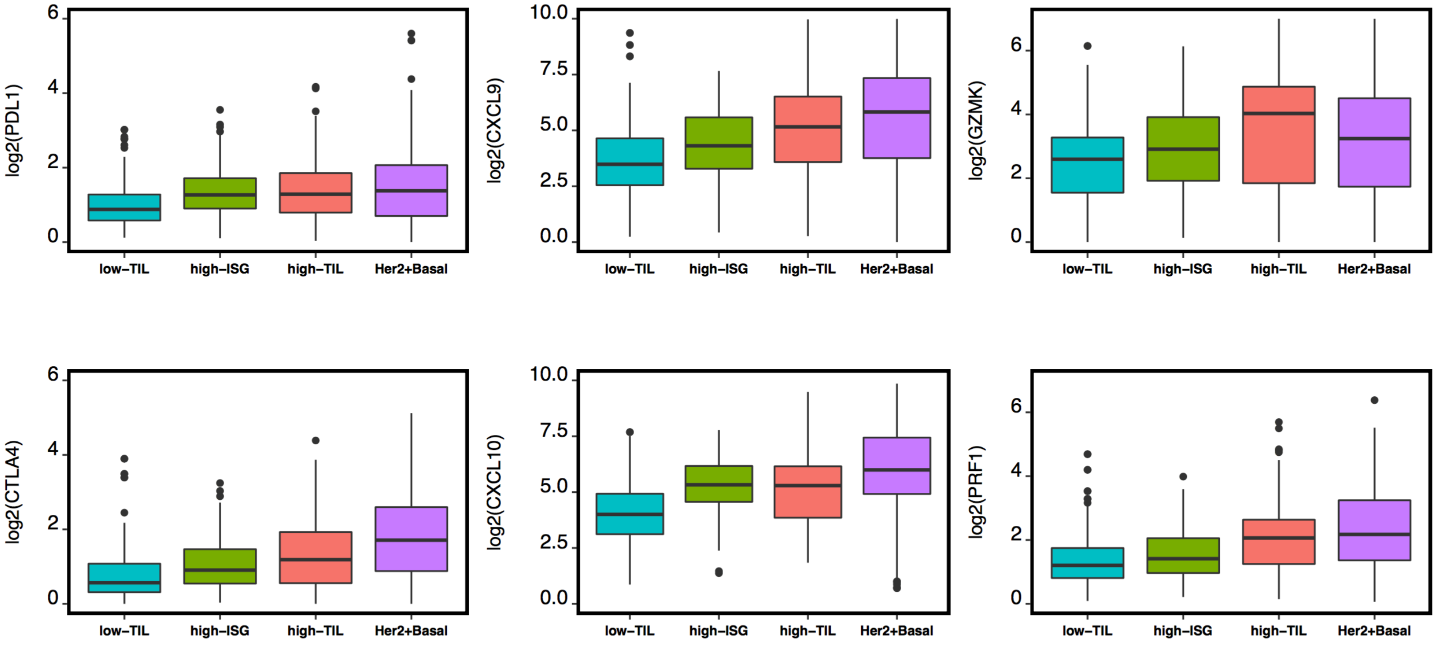


**Supplementary Figure 4.c**

### TCGA

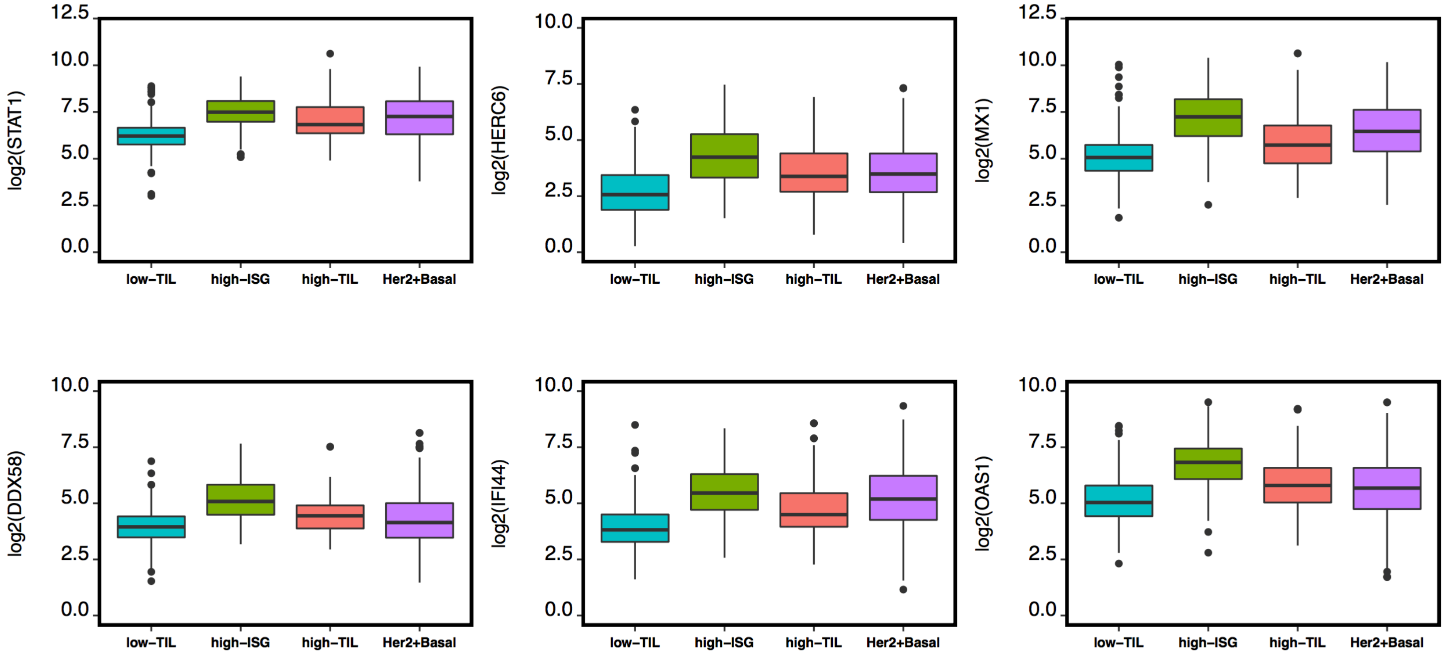


### KBC

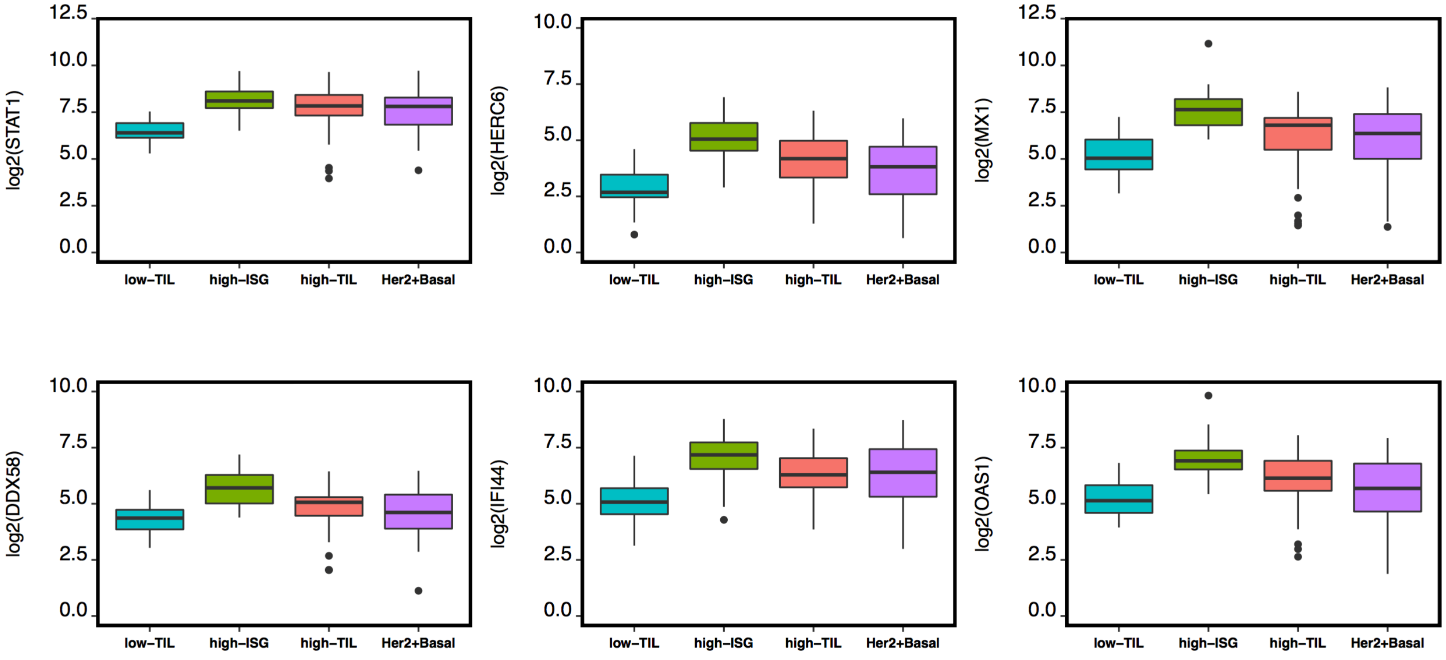


**Supplementary Figure 4.d**

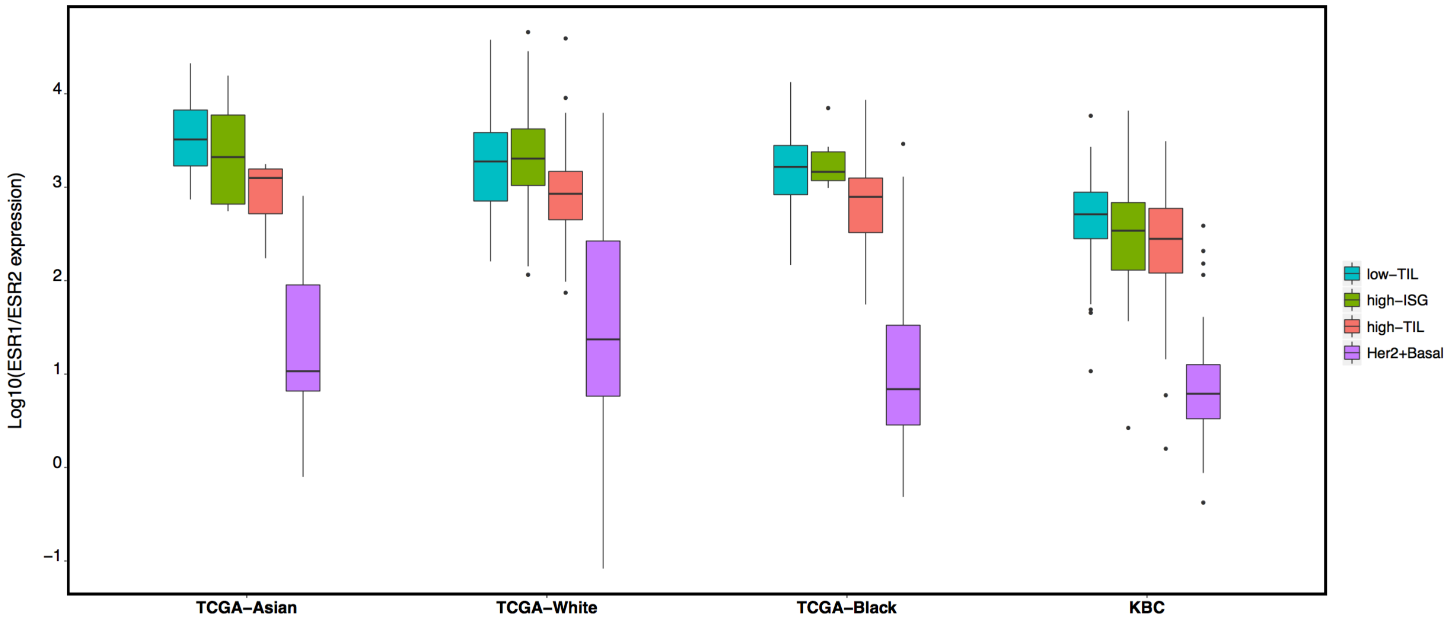


**Supplementary Figure 5.a**

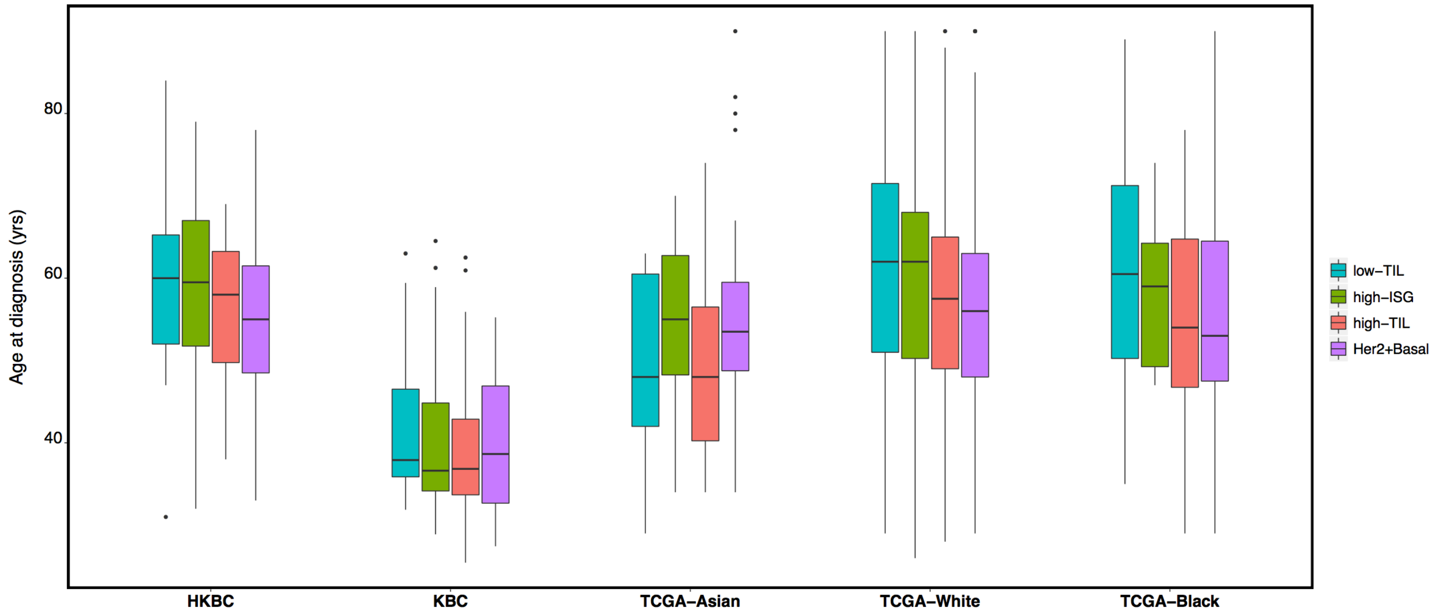


**Supplementary Figure 5.b**

### TCGA BRCA 10-year survival

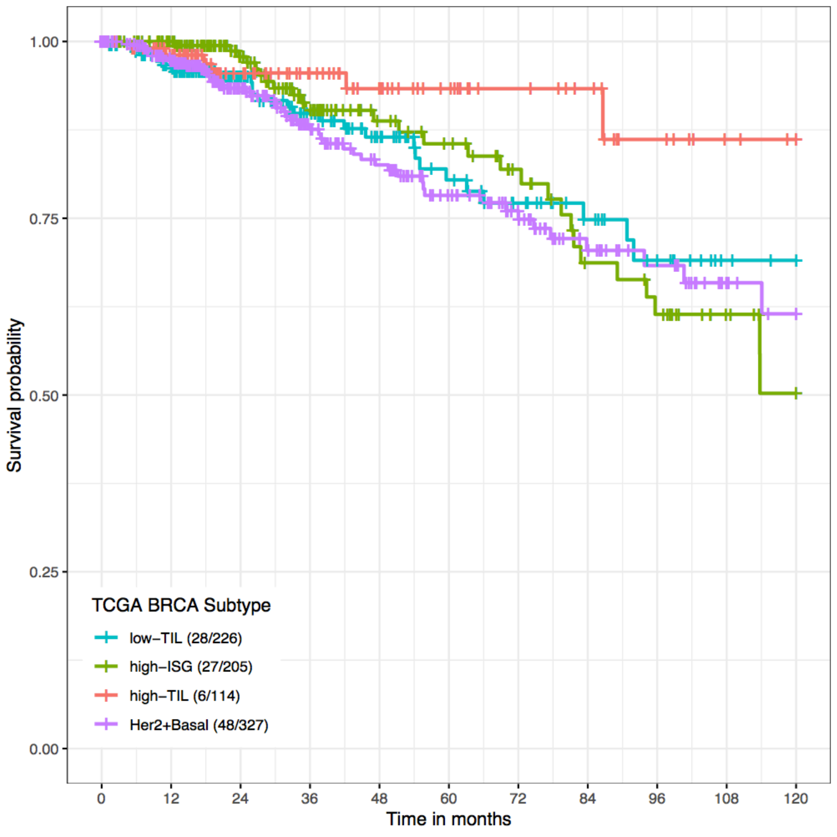


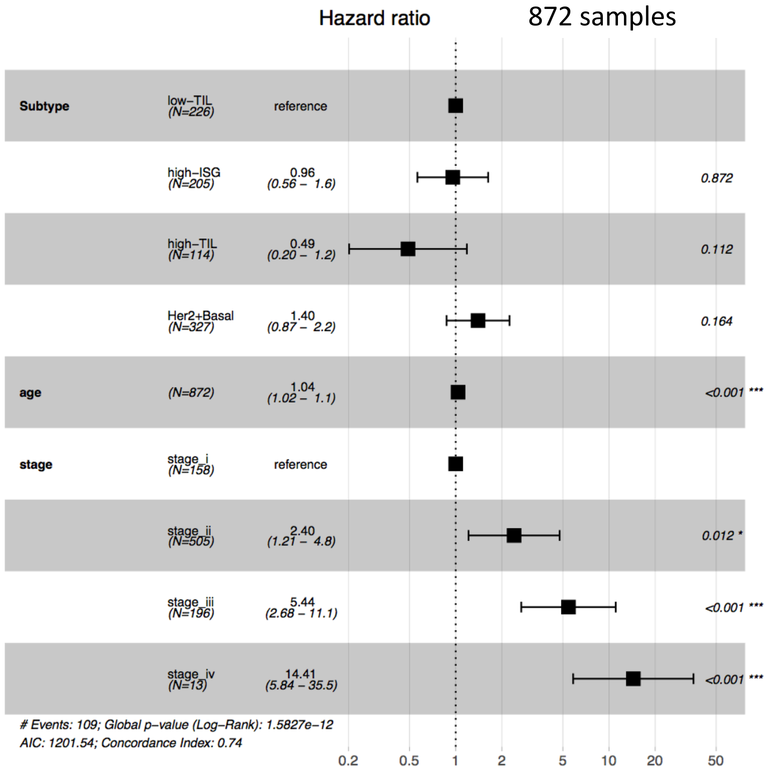


**Supplementary Figure 5.c**

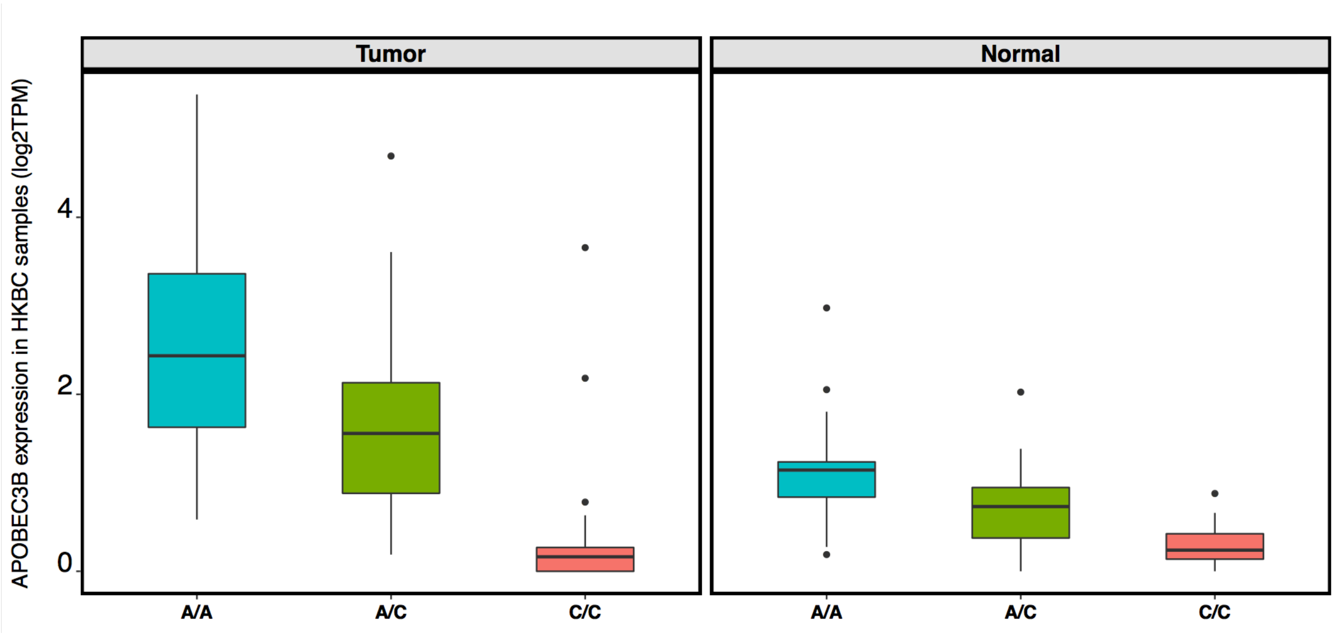


**Supplementary Figure 6**

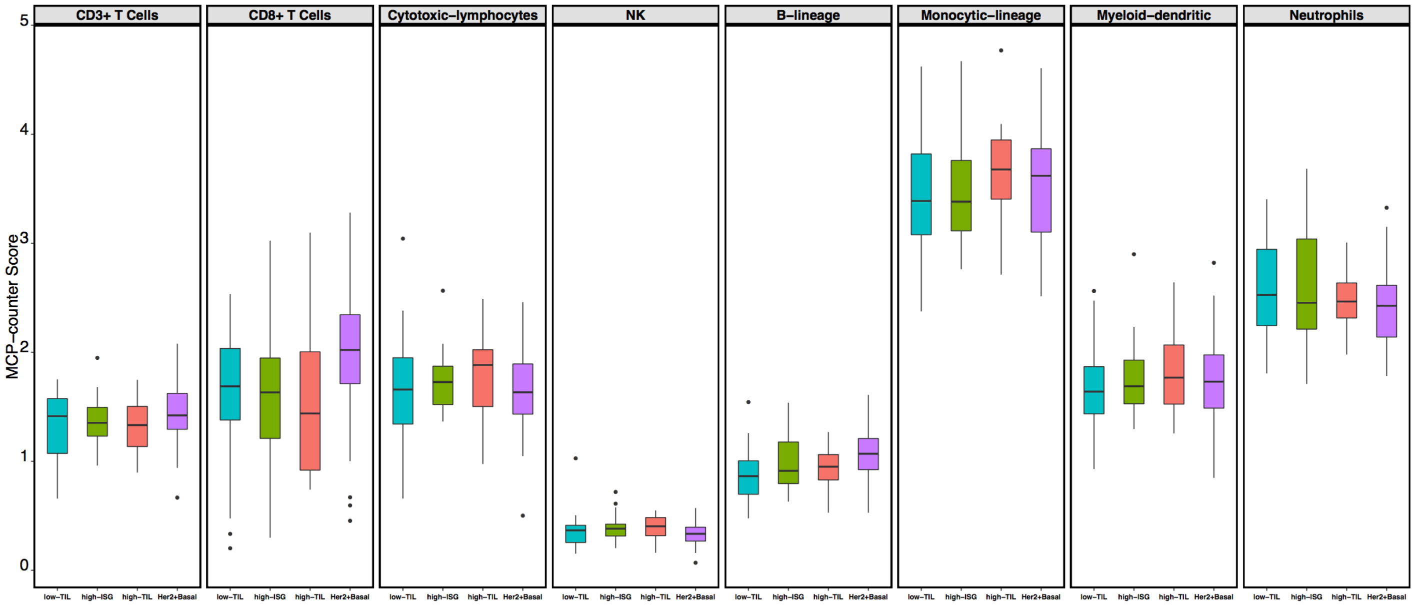


**Supplementary Figure 7**
