## Supplementary tables for "Immune gene expression profiling reveals heterogeneity in luminal breast tumors"

**Supplementary Table 1** - The distribution of clinical characteristics and key breast cancer risk factors in the Hong Kong breast cancer study (HKBC)

**Supplementary Table 2** - 130 immune-related genes used for the classification of luminal tumors

**Supplementary Table 3** - The three immune subtypes in relation to the luminal A/B classification in HKBC

**Supplementary Table 4** - P values for comparisons of tumor MCP-counter abundance scores by immune subtype in HKBC

**Supplementary Table 5** - P values for comparisons of tumor CIBERSORT fraction scores by immune subtype in HKBC

**Supplementary Table 6** - P values for comparisons of the MCP-counter abundance scores between paired tumor and normal tissue (N=80) by immune subtype in HKBC

**Supplementary Table 1. The distribution of clinical characteristics and key breast cancer risk factors in the Hong Kong breast cancer study (HKBC)**

| | Overall (n=139) | | low-TIL (n=40) | | high-ISG (n=36) | | high-TIL (n=16) | | |
|---|---|---|---|---|---|---|---|---|---|
| | N | % | N | % | N | % | N | % | p-value[a] |
| **Age at diagnosis (years)** | | | | | | | | | |
| **mean, SD** | 58.0 | 11.0 | 60.2 | 10.4 | 58.1 | 12.0 | 56.6 | 8.5 | 0.4688 |
| <50 | 31 | 22.3 | 6 | 15.0 | 8 | 22.2 | 4 | 25.0 | |
| 50-60 | 45 | 32.4 | 13 | 32.5 | 10 | 27.8 | 5 | 31.3 | 0.8918 |
| >60 | 63 | 45.3 | 21 | 52.5 | 18 | 50.0 | 7 | 43.8 | |
| **BMI** | | | | | | | | | |
| **mean, SD** | 24.8 | 4.0 | 24.1 | 3.8 | 24.6 | 4.4 | 27.9 | 3.7 | **0.0083** |
| <25 | 79 | 60.8 | 28 | 71.8 | 21 | 65.6 | 3 | 20.0 | |
| 25-30 | 33 | 25.4 | 7 | 17.9 | 7 | 21.9 | 8 | 53.3 | **0.0189** |
| >30 | 18 | 13.8 | 4 | 10.3 | 4 | 12.5 | 4 | 26.7 | |
| Missing | 9 | | 1 | | 4 | | 1 | | |
| **Age at menarche (years)** | | | | | | | | | |
| **mean, SD** | 13.8 | 2.1 | 13.9 | 2.2 | 13.6 | 1.9 | 13.6 | 2.0 | 0.8321 |
| <12 | 15 | 11.4 | 6 | 16.2 | 2 | 5.9 | 3 | 18.8 | |
| 12-14 | 70 | 53.0 | 17 | 45.9 | 20 | 58.8 | 8 | 50.0 | 0.6685 |
| >14 | 47 | 35.6 | 14 | 37.8 | 12 | 35.3 | 5 | 31.3 | |
| Missing | 7 | | 3 | | 2 | | | | |
| **Menopausal status** | | | | | | | | | |
| Premenopausal | 37 | 27.6 | 10 | 25.6 | 9 | 26.5 | 2 | 13.3 | 0.5739 |
| Postmenopausal | 97 | 72.4 | 29 | 74.4 | 25 | 73.5 | 13 | 86.7 | |
| Missing | 5 | | 1 | | 2 | | 1 | | |
| **Age at menopause (years)** | | | | | | | | | |
| **mean, SD** | 50.1 | 4.7 | 50.0 | 4.2 | 51.1 | 5.0 | 49.6 | 4.7 | 0.6425 |
| <50 | 25 | 30.9 | 7 | 30.4 | 4 | 19.0 | 5 | 50.0 | 0.4982 |
| >50 | 56 | 69.1 | 16 | 69.6 | 17 | 81.0 | 5 | 50.0 | |
| Missing | 58 | | 17 | | 15 | | 6 | | |
| **Parity** | | | | | | | | | |
| 0 | 16 | 12.2 | 2 | 5.3 | 8 | 23.5 | 3 | 21.4 | |
| 1-2 | 83 | 63.4 | 24 | 63.2 | 18 | 52.9 | 6 | 42.9 | 0.3169 |
| 3+ | 32 | 24.4 | 12 | 31.6 | 8 | 23.5 | 5 | 35.7 | |
| Missing | 8 | | 2 | | 2 | | 2 | | |
| **Age at first birth (years)** | | | | | | | | | |
| **mean, SD** | 26.8 | 4.9 | 26.8 | 5.3 | 26.5 | 4.2 | 25.5 | 5.1 | 0.7190 |
| <25 | 42 | 37.8 | 14 | 38.9 | 10 | 41.7 | 5 | 41.7 | |
| 25-30 | 38 | 34.2 | 12 | 33.3 | 9 | 37.5 | 5 | 41.7 | 0.3187 |
| >30 | 31 | 27.9 | 10 | 27.8 | 5 | 20.8 | 2 | 16.7 | |
| Missing | 28 | | 4 | | 12 | | 4 | | |
| **Grade** | | | | | | | | | |
| 1 | 11 | 12.2 | 3 | 12.5 | 4 | 18.2 | 2 | 22.2 | |
| 2 | 40 | 44.4 | 15 | 62.5 | 9 | 40.9 | 4 | 44.4 | 0.6481 |
| 3 | 39 | 43.3 | 6 | 25.0 | 9 | 40.9 | 3 | 33.3 | |
| Missing | 49 | | 16 | | 14 | | 7 | | |
| **Stage** | | | | | | | | | |
| I/II | 69 | 77.5 | 20 | 80.0 | 20 | 74.1 | 9 | 100.0 | 0.8777 |
| III | 20 | 22.5 | 5 | 20.0 | 7 | 25.9 | 0 | 0.0 | |
| Missing | 50 | | 15 | | 9 | | 7 | | |
| **Nodal Status** | | | | | | | | | |
| Present | 28 | 32.9 | 7 | 28.0 | 6 | 24.0 | 4 | 44.4 | 0.5060 |
| Absent | 57 | 67.1 | 18 | 72.0 | 19 | 76.0 | 5 | 55.6 | |
| Missing | 54 | | 15 | | 11 | | 7 | | |
| **Pam50 Subtype** | | | | | | | | | |
| Luminal A | 49 | 35.3 | 27 | 67.5 | 12 | 33.3 | 10 | 62.5 | **0.0084** |
| Luminal B | 43 | 30.9 | 13 | 32.5 | 24 | 66.7 | 6 | 37.5 | |
| Her2 | 20 | 14.4 | | | | | | | |
| Basal-like | 20 | 14.4 | N/A | | | | | | |
| Normal-like | 7 | 5.0 | | | | | | | |

low-TIL(Lum1), high-ISG(Lum2), high-TIL(Lum3)

[a] p-values were obtained using one-way analysis of variance (ANOVA) for continuous variables or chi-square tests for categorical variables

**anything in boldface represents a significant result**

### Supplementary Table 2: 130 Immune-related genes used for classification of luminal tumors

| | T cell signatures | | | Activated CD8/NK cells | Antigen presenting cells | NK cells | Macrophages | Presentation | Interferon-related genes | | T-cell regulation | |
|---|---|---|---|---|---|---|---|---|---|---|---|---|
| **Immune Score** | **LCK** | **Tfh** | **Tregs** | **Cytolytic** | **MHC2** | **NK** | **Macrophages** | **MHC1** | **STAT1** | **IF_I** | **Co_stimulation** | **Co_inhibition** |
| PRF1 | ARHGAP15 | CD200 | FOXP3 | GZMA | HLA-DMA | KLRF1 | FUCA1 | HLA-A | CXCL10 | DDX58 | CD2 | BTLA |
| GZMB | ARHGAP25 | CXCL13 | C15orf53 | PRF1 | HLA-DQB1 | KLRC1 | MMP9 | HLA-B | CXCL11 | HERC6 | CD226 | C10orf54 |
| CXCL13 | CCL5 | FBLN7 | IL5 | | HLA-DRA | | LGMN | HLA-C | GBP1 | IFI44 | CD27 | CD160 |
| IRF1 | CCR2 | ICOS | CTLA4 | | HLA-DRB4 | | HS3ST2 | HLA-F | STAT1 | IFI44L | CD28 | CD244 |
| IKZF1 | CCR7 | SGPP2 | IL32 | | | | TM4SF19 | HLA-G | | IFIT1 | CD40 | CD274 |
| HLA-E | CD2 | SH2D1A | GPR15 | | | | CLEC5A | HLA-J | | IFIT2 | CD40LG | CTLA4 |
| | CD247 | TIGIT | IL4 | | | | GPNMB | | | MX1 | CD58 | HAVCR2 |
| | CD27 | PDCD1 | | | | | C11orf45 | | | OAS1 | CD70 | LAG3 |
| | CD3D | | | | | | CD68 | | | OAS3 | ICOS | LAIR1 |
| | CD48 | | | | | | CYBB | | | RSAD2 | ICOSLG | LGALS9 |
| | CD53 | | | | | | | | | | SLAMF1 | PDCD1LG2 |
| | CORO1A | | | | | | | | | | TNFRSF18 | PVRL3 |
| | CSF2RB | | | | | | | | | | TNFRSF25 | TIGIT |
| | EVI2B | | | | | | | | | | TNFRSF4 | |
| | FGL2 | | | | | | | | | | TNFRSF8 | |
| | GIMAP4 | | | | | | | | | | TNFRSF9 | |
| | GIMAP5 | | | | | | | | | | TNFSF14 | |
| | GMFG | | | | | | | | | | TNFSF15 | |
| | GZMA | | | | | | | | | | TNFSF18 | |
| | GZMK | | | | | | | | | | TNFSF4 | |
| | HCLS1 | | | | | | | | | | TNFSF8 | |
| | IL10RA | | | | | | | | | | TNFSF9 | |
| | IL2RG | | | | | | | | | | | |
| | IL7R | | | | | | | | | | | |
| | INPP5D | | | | | | | | | | | |
| | IRF8 | | | | | | | | | | | |
| | ITK | | | | | | | | | | | |
| | KLRK1 | | | | | | | | | | | |
| | LCK | | | | | | | | | | | |
| | LCP2 | | | | | | | | | | | |
| | LPXN | | | | | | | | | | | |
| | LTB | | | | | | | | | | | |
| | PIK3CD | | | | | | | | | | | |
| | PLAC8 | | | | | | | | | | | |
| | PRG1 | | | | | | | | | | | |
| | PRKCB1 | | | | | | | | | | | |
| | PTPRC | | | | | | | | | | | |
| | RAC2 | | | | | | | | | | | |
| | SAMSSN1 | | | | | | | | | | | |
| | SCYA5 | | | | | | | | | | | |
| | SELL | | | | | | | | | | | |
| | SD2D1A | | | | | | | | | | | |
| | SLA | | | | | | | | | | | |
| | SLAMF1 | | | | | | | | | | | |
| | STAT4 | | | | | | | | | | | |
| | TNFRSF7 | | | | | | | | | | | |
| | TRBC1 | | | | | | | | | | | |

### Supplementary Table 3: The three immune subtypes in relation to the luminal A/B classification in HKBC

| | | Immune Subtypes (based on 13 mega-gene sets) | | |
|---|---|---|---|---|
| | | **low-TIL** | **high-ISG** | **high-TIL** |
| **Luminal A/B Class** | **Lum A** | 27 (55.10%) | 12 (24.49%) | 10 (20.41%) |
| | **Lum B** | 13 (30.23%) | 24 (55.81%) | 6 (13.95%) |

low-TIL(Lum1), high-ISG(Lum2), high-TIL(Lum3)

### Supplementary Table 4: Comparisons of tumor MCP-counter abundance scores by immune subtype in HKBC

| | low-TIL | high-TIL estimate | p-value[a] | high-ISG | high-TIL estimate | p-value[a] |
|---|---|---|---|---|---|---|
| **TCells** | Ref | 4.0971 | **<.0001** | Ref | 3.0447 | **0.0006** |
| **Tcells CD8** | | 3.4117 | **<.0001** | | 2.7796 | **0.0003** |
| **Cytotoxic Lymphocytes** | | 4.7760 | **<.0001** | | 3.6718 | **0.0005** |
| **NK Cells** | | 9.8510 | **0.0025** | | 7.1716 | **0.0118** |
| **B Lineage Cells** | | 2.3256 | **0.0002** | | 1.6216 | **0.0023** |
| **Monocytic Lineage** | | 1.0279 | **0.0411** | | 0.9784 | 0.0544 |
| **Myeloid Dendritic** | | 2.2858 | **0.0004** | | 2.9749 | **<.0001** |
| **Neutrophils** | | -0.7119 | 0.5736 | | 2.4914 | **0.0497** |

[a] p-values were obtained from age-adjusted logistic regression models

**anything in boldface represents a significant result**

### Supplementary Table 5: Comparisons of tumor CIBERSORT fraction scores by immune subtype in HKBC

| | low-TIL (n=40) | | high-ISG (n=36) | | high-TIL (n=16) | |
|---|---|---|---|---|---|---|
| | mean | SD | mean | SD | mean | SD |
| **Tcells CD8** | 0.0734 | 0.0582 | 0.0756 | 0.0413 | 0.1443 | 0.0302 |
| **Tcells CD4 memory resting** | 0.1656 | 0.0635 | 0.1730 | 0.0509 | 0.1715 | 0.0484 |
| **Tcells follicular helper** | 0.0702 | 0.0332 | 0.0950 | 0.0402 | 0.0839 | 0.0308 |
| **Monocytes** | 0.0591 | 0.0349 | 0.0497 | 0.0332 | 0.0491 | 0.0160 |
| **Macrophages M0** | 0.0585 | 0.0549 | 0.0499 | 0.0424 | 0.0194 | 0.0289 |
| **Macrophages M1** | 0.0536 | 0.0175 | 0.0697 | 0.0184 | 0.0725 | 0.0160 |
| **Macrophages M2** | 0.2519 | 0.0594 | 0.2280 | 0.0810 | 0.1939 | 0.0274 |

| | low-TIL | high-ISG | | high-TIL | |
|---|---|---|---|---|---|
| | | OR[a] (95% CI) | p-value | OR[a] (95% CI) | p-value |
| **Tcells CD8** | Ref | 1.00 (0.91-1.10) | 0.9906 | **1.44 (1.19-1.75)** | **0.0002** |
| **Tcells CD4 memory resting** | | 1.02 (0.94-1.10) | 0.5848 | 1.02 (0.91-1.13) | 0.7652 |
| **Tcells follicular helper** | | **1.23 (1.06-1.44)** | **0.0078** | 1.10 (0.92-1.33) | 0.3023 |
| **Monocytes** | | 0.92 (0.79-1.06) | 0.2382 | 0.91 (0.75-1.10) | 0.3561 |
| **Macrophages M0** | | 0.97 (0.88-1.06) | 0.4696 | **0.75 (0.61-0.93)** | **0.0095** |
| **Macrophages M1** | | **1.66 (1.25-2.22)** | **0.0005** | **1.84 (1.26-2.69)** | **0.0015** |
| **Macrophages M2** | | 0.95 (0.88-1.02) | 0.1673 | **0.86 (0.76-0.96)** | **0.0088** |

[a] p-values were obtained from age-adjusted logistic regression models

**anything in boldface represents a significant result**

### Supplementary Table 6: Comparisons of the MCP-counter abundance scores between paired tumor and normal tissue (N=80) by tumor subtype in HKBC

| | **low-TIL** | **high-ISG** | **high-TIL** | **non-Lum** |
|---|---|---|---|---|
| | | **p-value**[a] | | |
| **Tcells** | **0.0043** | 0.7481 | **0.0001** | **0.0036** |
| **Tcells CD8** | **0.0333** | 0.4177 | **0.0002** | **0.0253** |
| **Cytotoxic Lymphocytes** | **<.0001** | **<.0001** | 0.5306 | 0.6293 |
| **NK Cells** | **<.0001** | **<.0001** | **0.0309** | 0.7454 |
| **B Lineage Cells** | 0.1970 | 0.7567 | **0.0006** | **0.0146** |
| **Monocytic Lineage** | 0.3992 | 0.2590 | 0.6460 | 0.0796 |
| **Myeloid Dendritic** | **0.0105** | **<.0001** | 0.3519 | 0.3240 |
| **Neutrophils** | **<.0001** | **<.0001** | **0.0002** | **<.0001** |

[a] p-values were obtained from comparing tumor and normal samples using matched t-test

**anything in boldface represents a significant result**
